# Massively parallel characterization of CYP2C9 variant enzyme activity and abundance

**DOI:** 10.1101/2021.03.12.435209

**Authors:** Clara J. Amorosi, Melissa A. Chiasson, Matthew G. McDonald, Lai Hong Wong, Katherine A. Sitko, Gabriel Boyle, John P. Kowalski, Allan E. Rettie, Douglas M. Fowler, Maitreya J. Dunham

**Affiliations:** Department of Genome Sciences, University of Washington, Seattle, WA; Department of Medicinal Chemistry, University of Washington, Seattle, WA; Department of Bioengineering, University of Washington, Seattle, WA

## Abstract

*CYP2C9* encodes a cytochrome P450 enzyme responsible for metabolizing up to 15% of small molecule drugs, and *CYP2C9* variants can alter the safety and efficacy of these therapeutics. In particular, the anti-coagulant warfarin is prescribed to over 15 million people annually and polymorphisms in *CYP2C9* can affect patient response leading to an increased risk of hemorrhage. We developed Click-seq, a pooled yeast-based activity assay to test thousands of variants. Using Click-seq, we measured the activity of 6,142 missense variants expressed in yeast. We also measured the steady-state cellular abundance of 6,370 missense variants expressed in a human cell line using Variant Abundance by Massively Parallel sequencing (VAMP-seq). These data revealed that almost two-thirds of CYP2C9 variants showed decreased activity, and that protein abundance accounted for half of the variation in CYP2C9 function. We also measured activity scores for 319 previously unannotated human variants, many of which may have clinical relevance.

## INTRODUCTION

Recent sequencing efforts have resulted in an avalanche of new variants, many of which are variants of uncertain significance (VUS) - variants identified through genetic testing whose functional significance is unknown. VUS hamper the implementation of precision medicine as they must be classified as pathogenic or benign before they can be used to inform clinical decisions. Over half of the missense variants in ClinVar^1^ are VUS^2^. VUS are a particular problem in the field of pharmacogenomics, which seeks to understand the genetic sources of inter-individual variation in drug response. Functionally annotated pharmacogene variants can be used to guide dosing decisions and predict adverse drug reactions (ADRs), which cost U.S. hospitals up to 30 billion dollars annually and are a leading cause of hospitalization and death^3, 4^. 30% of ADRs are predicted to be caused by inter-individual variability in drug metabolizing enzymes and other drug related genes^5^. Genetic variants predict drug response for a subset of important drugs, and implementing genotype-guided drug dosing can improve patient outcomes^6^. However, the vast majority of pharmacogene variants discovered so far are of unclear functional effect.

One important group of pharmacogenes is the Cytochromes P450 (CYPs). CYPs are a superfamily of monooxygenase enzymes that use heme as a cofactor, and there are 57 CYP genes in humans^7^. *CYP2C9* in particular is the primary metabolic enzyme for a wide range of drugs including drugs that must be dosed carefully such as phenytoin (for seizures) and the widely prescribed oral anticoagulant warfarin^8, 9^. *CYP2C9* polymorphisms contribute to an estimated 15% of the variation in warfarin dose^10^, and some common coding variants have large effects. For example, the *CYP2C9* I359L missense allele results in substantially diminished S-warfarin clearance leading to warfarin sensitivity^11^. Genotype-guided warfarin dosing based on *CYP2C9* and *VKOR* alleles can improve patient treatment in some situations^12^, but relies on knowing the function of alleles to guide dosing decisions.

Only a subset of CYP alleles have been studied adequately for genotype-guided dosing. As human CYP alleles are discovered, they are named according to the star (*) system^13^ and curated by the PharmVar consortium^14^. There are 70 documented *CYP2C9* star alleles in the PharmVar database (pharmvar.org). The Clinical Pharmacogenetics Implementation Consortium (CPIC) reviews *in vitro* and *in vivo* evidence and provides clinical functional recommendations for *CYP2C9* and other pharmacogenes^15^. CPIC has provided clinical allele functional annotations for 36 of the 70 *CYP2C9* star alleles^16, 17^. However, there are many more *CYP2C9* alleles than those documented in PharmVar. *CYP2C9* has 8 common alleles (MAF > 1%)^18^ and hundreds of documented rare alleles (MAF < 1%)^19, 20^. In the population database gnomAD^20^, there are 466 missense alleles in *CYP2C9*, half of which are singletons. The vast majority of variation in *CYP2C9* is unannotated, and so knowing the functional consequence of existing and yet-to-be discovered alleles will help improve dosing of drugs cleared by CYP2C9. Thus, there is a need for a large-scale experimental effort to comprehensively characterize *CYP2C9* variants.

We used deep mutational scanning (DMS) to measure the enzyme activity and steady-state cellular abundance of thousands of CYP2C9 missense variants. DMS is a high-throughput method for probing variant function by applying a functional selection, enriching variants with high function and depleting variants with low function^21^. High-throughput DNA sequencing is then used to quantify the change in each variant’s frequency during the selection, yielding a functional score for every variant in the library. Selections can take many forms, but often couple variant function to cell growth or measure protein or ligand binding, and rarely measure enzyme activity directly (e.g. ^22^). DMS approaches have the potential to transform pharmacogenomic implementation^23, 24^, but so far have been applied to only a handful of pharmacogenes, including *TPMT*, *NUDT15*, and *VKORC1*^25–27^. No multiplexed method for quantifying enzyme activity directly in cells exists currently, precluding the quantification of variant effects on human CYP enzyme despite the clear need for such comprehensive functional data.

To meet this challenge, we developed Click-seq, a multiplexed, sequencing-based method for quantifying protein variant activity, and used it to measure the activity of 6,142 CYP2C9 missense variants expressed heterologously in yeast cells. Additionally, we leveraged the Variant Abundance by Massively Parallel sequencing assay we developed previously (VAMP-seq)^25^, which uses a fluorescent protein reporter coupled with FACS, to measure the abundance of 6,370 CYP2C9 missense variants expressed in cultured human cells. Comparison of both activity and abundance revealed that the mechanism behind variant loss of function could be attributed to reduced abundance for at least 50% of variants. Additionally, these data highlighted key regions of CYP2C9 crucial to function, including many residues involved in heme binding. Finally, our experimental functional scores are concordant with existing CYP2C9 functional annotations. We used our activity scores to annotate 319 previously unannotated human CYP2C9 missense variants in gnomAD, two-thirds of which had reduced activity. In addition to annotating these 319 variants, we provide activity scores for 5,797 additional variants. This information will be of great utility to clinicians as a key source of evidence when presented with VUS and will aid in improving dosing efficacy of drugs metabolized by CYP2C9.

## RESULTS

### Click-seq, a multiplexed assay for CYP2C9 enzymatic activity

We developed a multiplexed assay of CYP activity, Click-seq that uses a CYP-selective, activity-based probe to modify CYP variant enzymes heterologously expressed in the budding yeast *S. cerevisiae.* Following probe attachment via mechanism-based adduction, click chemistry is used to label the enzyme-bound probe with a fluorophore, FACS separates cells according to their degree of labeling, and high-throughput sequencing of the sorted cells is used to score each variant (Figure 1a). Click-seq directly measures enzyme activity by quantifying the amount of mechanism-based inhibitor covalently attached to the CYP enzymes in a cell after a period of incubation; thus labeling is activity-dependent. CYP-specific activity-based probes have been developed previously^28, 29^, but prior work has focused on *in vitro* assays (commonly CYP-rich microsomal preparations), rather than cell-based methods. We modified existing assays to work with intact yeast cells in a pooled format. We also synthesized a new activity-based probe, tienilic acid hexynyl amide (TAHA), that is an analog of tienilic acid, a known covalent inhibitor of CYP2C9^30^. TAHA showed better labeling than a generic P450 probe^29^ (Supplementary Figure 1). Additionally, to improve recombinant CYP activity, human P450 accessory proteins were integrated into a modified laboratory strain (see Methods) resulting in a humanized yeast strain.

**Figure 1.**
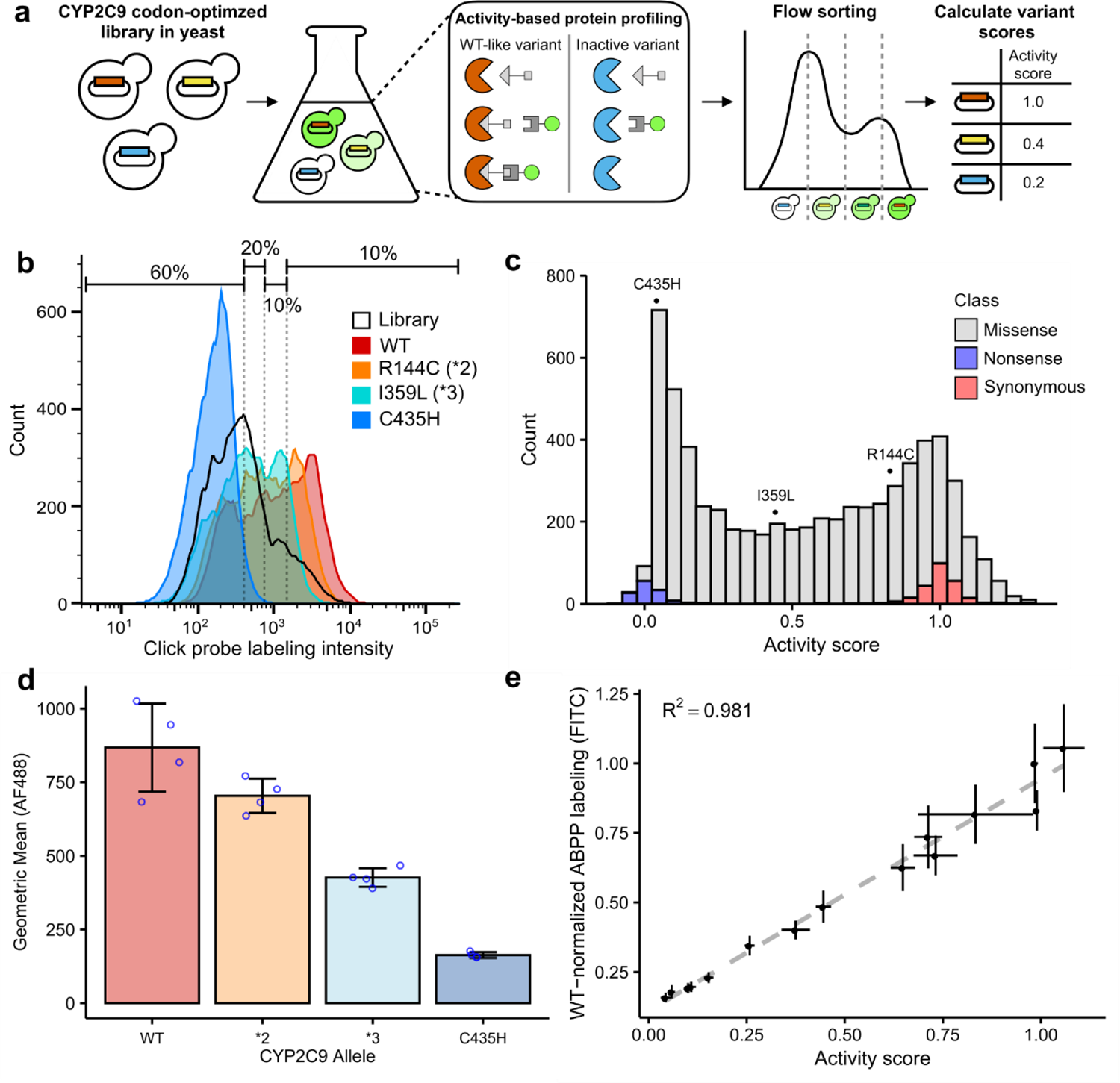
Multiplexed measurement of CYP2C9 activity using Click-seq. (a) A humanized yeast strain is transformed with a library of codon optimized *CYP2C9* variants, labeled using activity-based protein profiling (ABPP), resulting in a range of fluorescence levels, and sorted into four bins using fluorescence-activated cell sorting. Bins are sequenced to calculate relative variant activity. (b) Flow cytometry of ABPP labeled yeast expressing CYP2C9 WT (red), reduced activity alleles (*2 and *3, orange and turquoise), null allele (C435H, blue) and CYP2C9 variant library (black outline). Smoothed histograms shown, each sample represents ∼20,000 cells. Note that some cells with low intensity are the result of plasmid loss and thus do not contribute to the downstream sequencing results. (c) Stacked histogram of activity score colored by type of variant. Individual scores of C435H, *2, and *3 shown on top. (d) Geometric mean of ABPP-labeled CYP2C9 alleles. Individual replicates shown as blue points, and error bars show standard deviation. (e) WT-normalized ABPP labeling (FITC normalized fluorescence) for 14 CYP2C9 variants, expressed in the humanized yeast strain and labeled separately. Individual variants were labeled using the same ABPP protocol as the pooled assay. Scatter plot and linear regression of activity score (pool score) versus individual variant ABPP labeling (n = 3 replicates). Error bars show standard error for activity scores and standard error for ABPP labeling.

In order to demonstrate that Click-seq accurately reflects enzyme activity, we cloned individual CYP2C9 variants of known activity and compared probe labeling levels to wild type CYP2C9. We found that, as expected, CYP2C9 *2 (R144C) and *3 (I359L) had decreasing levels of labeling, and a catalytically inactive variant, C435H, had labeling comparable to background levels (Figure 1b, d). We then constructed a barcoded, site-saturation mutagenesis library of *CYP2C9* codon optimized for yeast expression and encompassing positions 2 to 490. This library covers 6,542 of the 9,780 possible single amino acid variants (67%), with 105,372 barcodes (mean of 5.8 and median of 3 for single amino acid variants; see Supplementary Table 3 for details). The CYP2C9 activity library was labeled using the TAHA probe and flow sorted into bins; DNA collected from each bin was amplified, sequenced, and analyzed to determine relative variant activity. We calculated activity scores (see Methods) for 6,524 single variants, of which 6,142 were missense, 131 were nonsense, and 250 were synonymous (Figure 1c). Activity scores were normalized to median nonsense and synonymous variant scores such that a score of 0 represented nonsense-like activity and a score of 1 represented wild-type-like activity. Variant activity scores correlated very well between the four replicate sorts we performed from four separate library outgrowths (mean Pearson’s *r* = 0.92, mean Spearman’s *ρ* = 0.919, Supplementary Figure 3). We binned activity scores into activity classes (Methods and Supplementary Figure 4) and found that a 64.9% (3,987) of missense variants showed significantly decreased activity compared to wild type. As further confirmation that our classifications align with existing standards, the boundary between “WT-like” activity and “decreased” activity is very close to the activity score for CYP2C9 *2, a known decreased activity allele.

To internally validate our Click-seq derived activity data, we generated 14 CYP2C9 variants that spanned the full range of activity scores, labeled them individually, and found that individually tested and Click-seq derived activity scores were well correlated (Pearson’s *r* = 0.991, Figure 1e). To show that our large-scale activity scores determined with the TAHA probe were representative of CYP2C9 variant activity towards important CYP2C9 drug substrates, we performed gold-standard LC-MS assays of S-warfarin 7-hydroxylation and phenytoin 4-hydroxylation using microsomal preparations derived from yeast expressing the same 14 CYP2C9 variants generated for internal validation (Figure 2). Activity scores were well correlated with individual variant S-warfarin turnover (Pearson’s *r* = 0.874, Spearman’s *ρ* = 0.895) and phenytoin turnover (Pearson’s *r* = 0.764, Spearman’s *ρ* = 0.87). Both of these CYP2C9 drug substrates had highly similar activity levels across the variants tested (Pearson’s *r* = 0.965, Spearman’s *ρ* = 0.979, Supplementary Figure 5). Additionally, we found that individual variant activity scores correlated well with an assay based on a fluorogenic substrate, BOMCC (Supplementary Figure 6), indicating consistency across methods, as fluorogenic substrate assays are another standard method of measuring CYP activity^31^.

**Figure 2.**
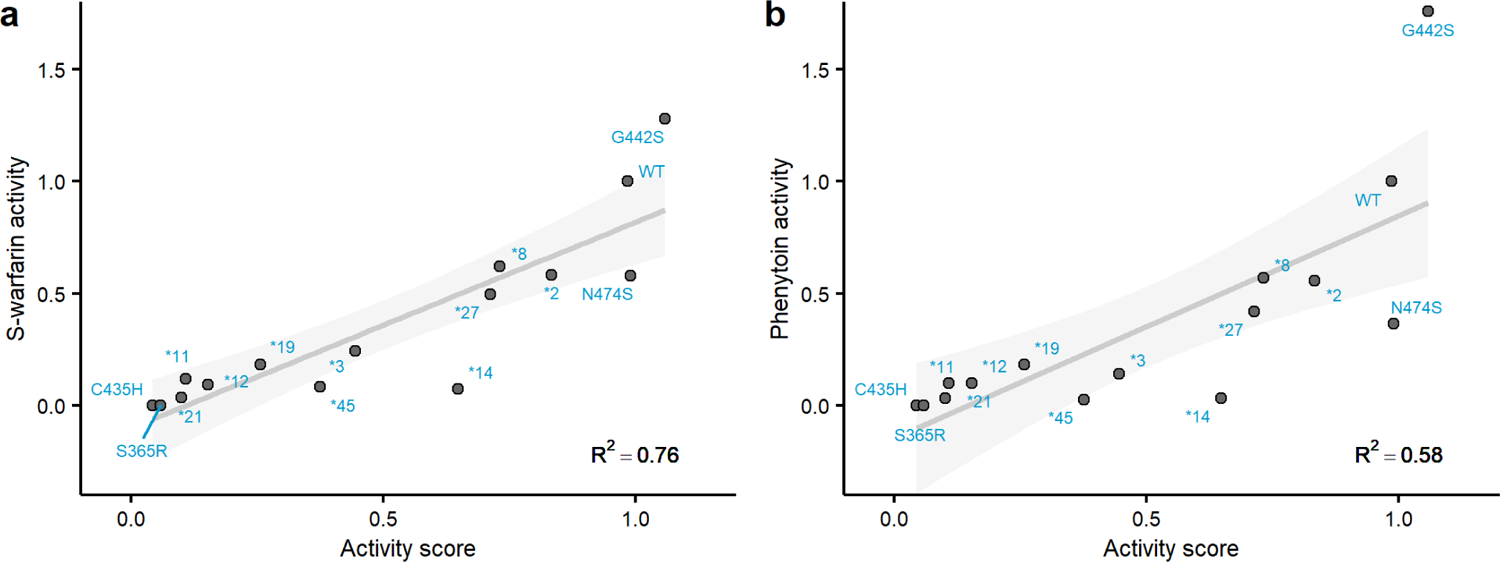
Comparison of CYP2C9 activity scores with gold-standard activity assays on yeast microsomes. Scatterplots of CYP2C9 activity scores plotted against individually tested CYP2C9 alleles. Individual alleles were expressed in the humanized yeast strain used in the pooled assay, and yeast microsomes were harvested from these individual strains. In a), LC-MS was used to determine the rate of S-warfarin 7-hydroxylation. In b), LC-MS was used to determine the rate of phenytoin 4-hydroxylation. The grey line is the regression line, and shaded area shows the 95% confidence interval. All activities are shown normalized to wild type rates.

Overall, Click-seq yielded a map relating variant sequence to activity, but did not provide information on the mechanisms underlying variant loss of function. Thus, we performed a second CYP2C9 DMS, scoring variants for their abundance in cells in order to determine to what degree decreases in variant activity could be explained by decreases in abundance.

### A multiplexed assay for CYP2C9 abundance in cultured human cells

We recently developed a method, VAMP-seq^25^, that enables measurement of steady-state protein abundance in cultured human cell lines using fluorescent reporters (Figure 3a). We applied VAMP-seq to CYP2C9, fusing eGFP C-terminally (Supplementary Figure 7), and from the same construct expressing mCherry via an internal ribosomal entry site (IRES) to control for cell-to-cell differences in expression. The fluorescent reporters accurately quantified the loss of abundance of a known destabilized CYP2C9 variant^32^, R335W (*11), relative to wild type as measured by the ratio of eGFP to mCherry (Figure 3b). We constructed a barcoded, site-saturation mutagenesis library of CYP2C9, encompassing positions 2 to 490. This library covered 8,310 of the 9,780 possible single amino acid variants (85%), with 78,740 barcodes (mean of 5.9 and median of 4 for single amino acid variants; see Supplementary Table 3 for details).

**Figure 3.**
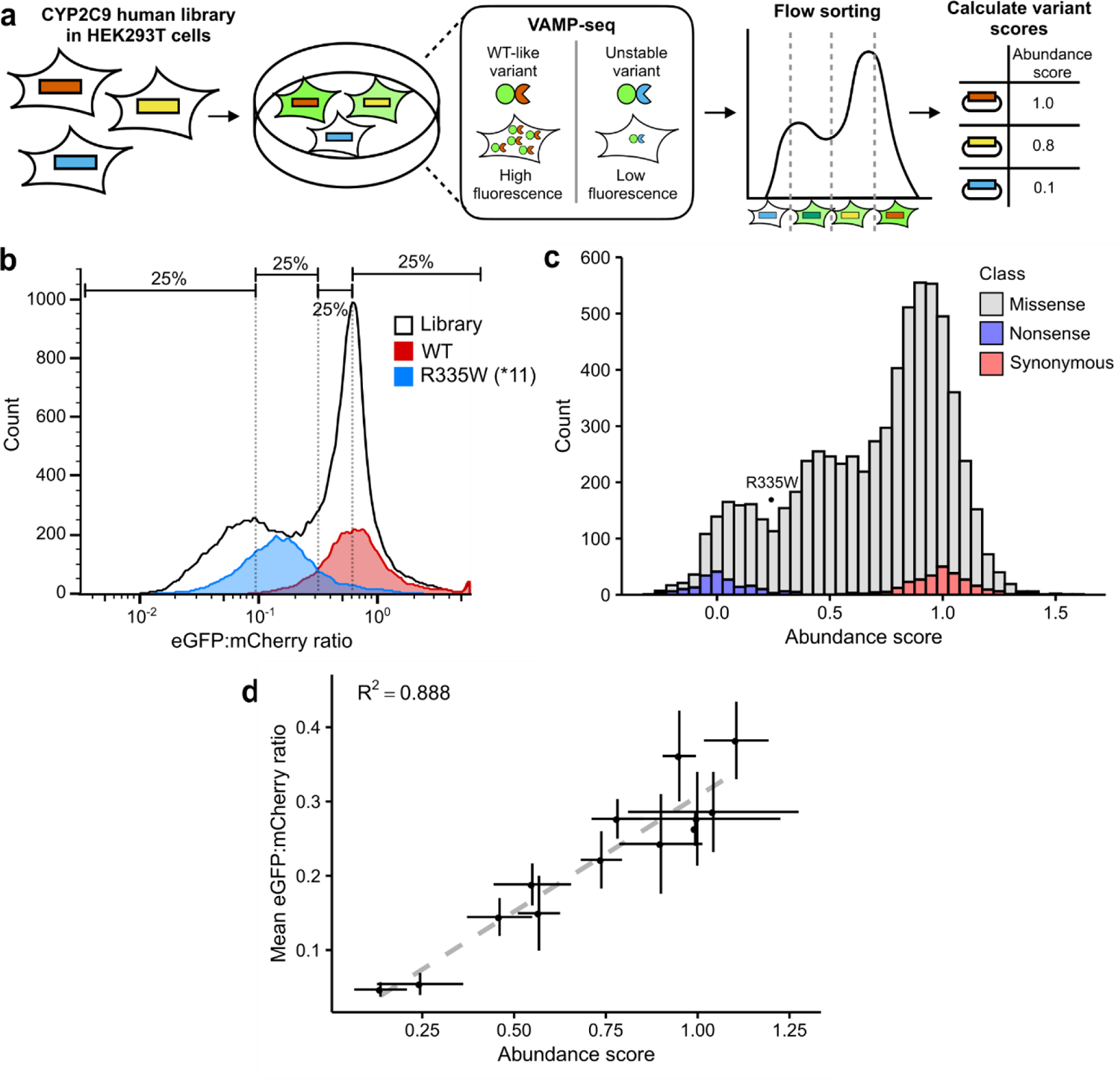
Multiplexed measurement of CYP2C9 activity using VAMP-seq. (a) Using VAMP-seq^25^, a *CYP2C9* library was expressed in HEK 293T cells such that each variant was expressed as an eGFP fusion, resulting in a range of fluorescence according to variant stability. Cells were then flow sorted into bins and sequenced to determine relative variant abundance. (b) Flow cytometry of CYP2C9 WT (red), destabilizing allele (*11, blue), and CYP2C9 eGFP fusion library expressed in HEK293T cells (black outline). Smoothed histograms of eGFP:mCherry ratios shown. Approximate quartile bins for sorting shown at the top. (c) Stacked histogram of abundance score colored by type of variant. Abundance score of *11 shown as a point. (d) Scatterplot and linear regression of individually measured cell eGFP:mCherry ratios for 12 CYP2C9 variants vs VAMP-seq derived abundance scores for the same variants. Error bars show standard error for abundance scores and standard error for individually determined eGFP:mCherry ratio (n = 2 replicates).

We expressed this library in HEK 293T cells using a serine integrase landing pad^25, 33^. Successfully recombined cells expressing CYP2C9 variants were selected with a small molecule, AP1903, and then sorted into quartile bins based on eGFP:mCherry ratio (Figure 3a). Bins were deeply sequenced, and the resulting sequencing reads were used to calculate frequencies across bins for each variant. Abundance scores were calculated using weighted averages of variant frequencies and normalized to the scores of synonymous and nonsense variants as for the activity scores (see Methods). Variant abundance scores showed distinct, separable distributions of synonymous and nonsense variants, with missense variants spanning the range between them (Figure 3c). After filtering, we assigned variant scores to 6,821 single variants, of which 6,370 were missense, 189 were nonsense, and 261 were synonymous. Three replicate sorts from two separate transfections were performed on this library and the replicates correlated well (mean Pearson’s *r* = 0.789, mean Spearman’s *ρ* = 0.754, Supplementary Figure 3). To internally validate our VAMP-seq derived abundance data, we generated 12 CYP2C9 variants, expressed them individually, and found that individually measured and VAMP-seq derived abundance scores were well correlated (Pearson’s *r* = 0.942, Figure 3d). In contrast to the activity classes, only 36.8% of missense variants (2,347 variants) had a significantly decreased abundance class. This fraction is similar to other VAMP-seq studies of pharmacogene abundance, as 34% of VKOR missense variants showed significantly decreased abundance^27^.

### Mechanism of CYP2C9 variant loss of function

Between the Click-seq and VAMP-seq datasets, 8,091 missense variants had at least one functional score, and 4,421 variants had both activity and abundance scores (Figure 4). Among these variants, activity and abundance were strongly correlated (Pearson’s *r* = 0.748, Spearman’s *ρ* = 0.749) (Figure 4f). We observed an abundance threshold at a score of ∼0.5, below which variants had very low activity (median activity score ∼0.098), suggesting that for variants with abundance below this level, differences in Click-seq signal are too small to detect. Conversely, variants with abundance scores greater than 0.5 had a wider range of activity scores. The overall positive trend between abundance and activity scores revealed that 1) using engineered yeast as a heterologous CYP expression system largely recapitulates protein behavior in human cells, and 2) a substantial number of variants had low activity because they were less abundant. We estimated that approximately half of the variation in activity could be explained by abundance (*R^2^* = 0.56, Figure 4f). Since there was no normalization to protein expression per cell, the yeast activity scores reported are each a combination of both variant activity and variant stability.

**Figure 4.**
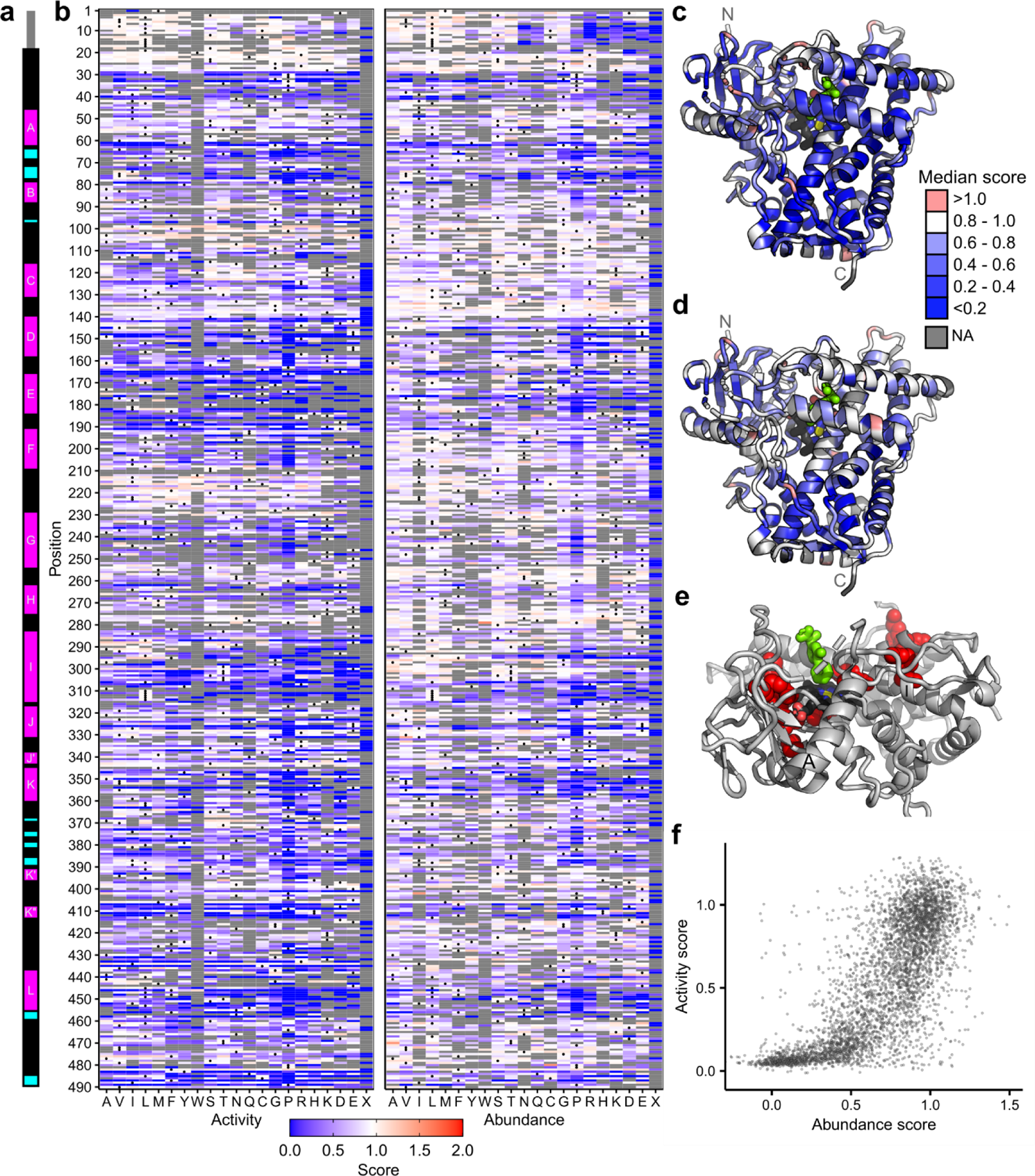
Click-seq activity scores and VAMP-seq abundance scores for CYP2C9. (a) Secondary structure of CYP2C9, with alpha helices in magenta and beta sheets in cyan. Helix names labeled in white. (b) Heatmaps of CYP2C9 activity (left) and abundance (right) scores. WT amino acids denoted with a dot, and missing data are shown in grey. Scores range from nonfunctional (blue) to WT-like (white) to increased (red). In (c) and (d), CYP2C9 structure (PDB: 1r9o) colored by median activity and abundance, respectively, at each position. Median scores are binned as depicted in legend, and missing positions shown in grey. Heme colored by element (carbon:black, nitrogen:blue, oxygen:red, iron:yellow), and substrate (flurbiprofen) colored bright green. Median activity scores shown in (c) and median abundance scores shown in (d). (e) Zoomed view of partial CYP2C9 structure. Positions with the lowest 2.5% specific activity scores shown as red spheres. F and G helices hidden, A and I helices labeled, and heme and substrate colored as in (c) and (d). (f) Scatter plot of CYP2C9 activity and abundance scores from a total of 4,421 missense variants.

By comparing activity and abundance, we were able to identify variants that abolish activity but not abundance. We hypothesized that functionally important regions such as the active site and binding pocket of CYP2C9 would be enriched for such low activity, high abundance variants. To find such variants, we calculated variant specific activity by dividing the activity score by the abundance score (see Methods). We found that the positions with the lowest median specific activity were not active site positions, but instead mainly positions likely involved in heme binding (Figure 4e). This finding implies that these positions are crucial for activity but did not strongly destabilize the protein when mutated. The positions with the lowest 2.5% median specific activity scores included the heme-binding motif residues Gly431, Arg433, Cys435, and Gly437^34^ (Supplementary Figure 8) as well as Arg97, which is important for heme propionate binding in CYP2C9^35^. Finally, we found that variants in the active site^36^ had median activity scores of 0.61 and median abundance scores of 0.91, indicating that these variants were generally not destabilizing and also only had moderate effects on activity. This active site mutational tolerance is surprising, at least in contrast to VKOR, where active site positions had the lowest specific activity scores^27^. However, CYPs have well-documented conformational flexibility, especially in the active site^37^. Moreover, in CYP2C9 the BC loop which frames the substrate access channel is also highly flexible^38^, with median activity and abundance scores of 0.81 and 0.88 respectively. Thus, CYP2C9 is apparently able to tolerate active site mutations without loss of abundance.

### Structural insights from CYP2C9 functional scores

ER-localized CYP enzymes are composed of an N-terminal ER-transmembrane domain and a large, cytoplasmic catalytic domain^39^. The CYP enzyme superfamily is diverse at the sequence level but members share a common structure including 12 major helices, labeled A through L, and four beta sheets, labeled β1 through β4^40^ (Figure 4a). CYP2C9 has been crystallized with warfarin^41^ and flurbiprofen^42^ as well as with other substrates^43^, and also without a ligand^41^. These structures are generally comparable and show small differences in substrate-interacting regions. We used the flurbiprofen-bound CYP2C9 structure for our analysis as the substrate is bound in a catalytically favorable orientation in this structure. Three positions are almost completely conserved across all CYPs: Glu354 and Arg357 in the ExxR motif involved in heme binding and core-stabilizing^44^, and also the invariant heme-coordinating cysteine, Cys435^45^. In our activity assay, the 36 missense variants at these three positions all had activity scores of < 0.1. In our abundance assay, Arg357 was also extremely intolerant to substitution with a median abundance score of 0.077, indicating that Arg357 is crucial for both activity and abundance. In addition to these highly conserved residues, our activity scores recapitulated the importance of the heme binding motif at positions 428 - 437^34^, and the proline-rich PPGP motif in the linker (or hinge) region after the transmembrane domain^46^ which is necessary for proper folding^47^ (Supplementary Figure 8).

We mapped median positional activity and abundance scores onto the CYP2C9 structure (Figure 4c,d) to identify key regions important for activity and abundance. To determine the characteristic mutation patterns in different regions of CYP2C9, we performed hierarchical clustering of positions based on both activity and abundance scores and identified six main clusters of positions (Figure 5). We found that substitutions in Cluster 3 were universally not tolerated, and positions in this Cluster generally grouped into two distinct regions: core-facing positions in helices D, E, I, J, K, and L comprising the highly conserved heme binding structural core of the protein, and positions in and directly abutting β sheet 1. Both of these regions are highly conserved across CYPs and are composed of buried, hydrophobic residues^44, 48^, in which substitution leads to destabilization and degradation. In addition, substitutions in β sheet 1 may disrupt distal side chains that coordinate with the central heme iron^49^. Clusters 1 and 2 were slightly more tolerant to substitution than Cluster 3 and are also found in the core of the protein.

**Figure 5.**
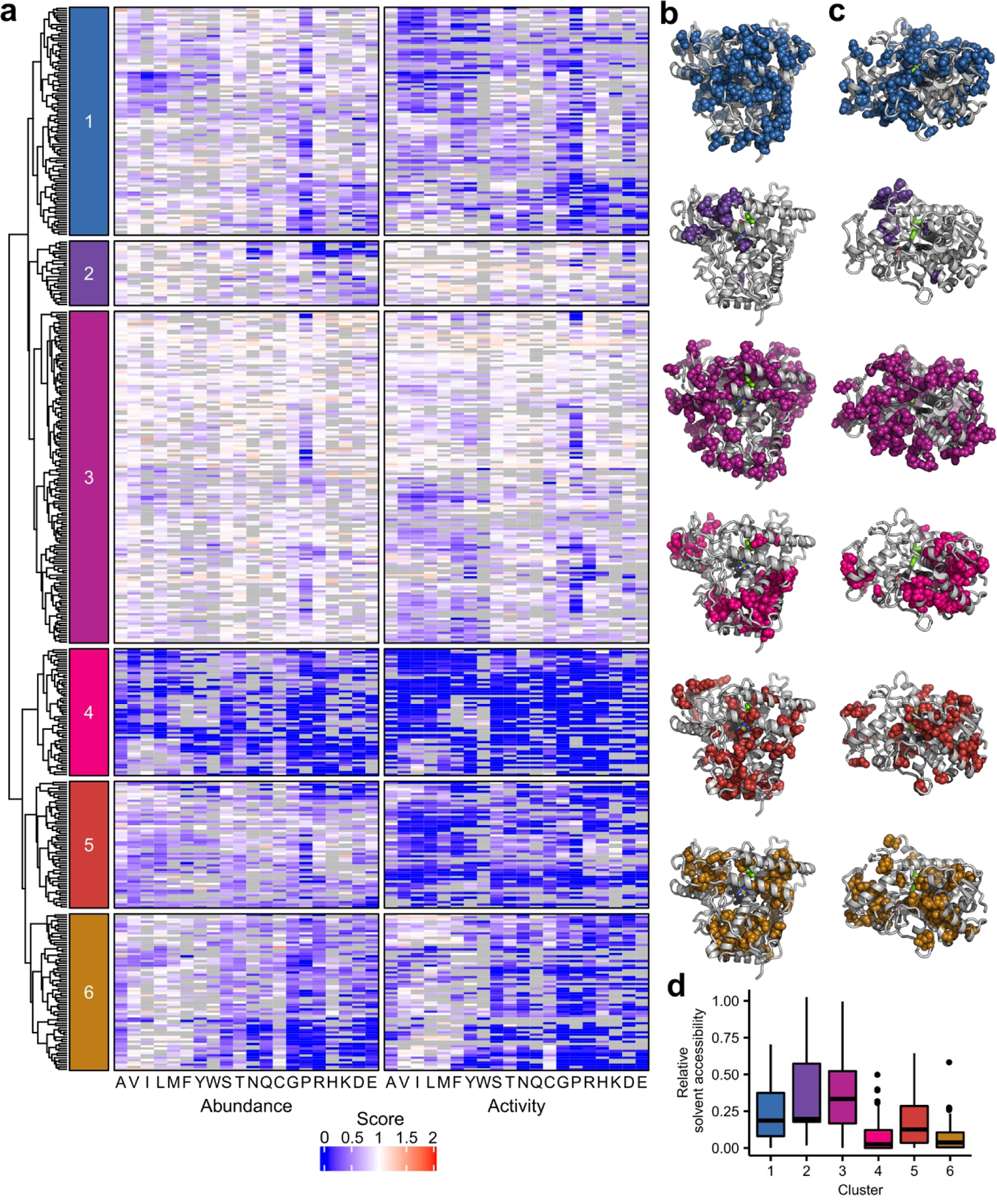
Hierarchical clustering of activity and abundance scores and cluster accessibility. In (a), dendrogram and heatmaps of CYP2C9 activity and abundance score clustered by position. Heatmaps colored as in Figure 4. Only positions that had at least 26 total mutations were included in this analysis. Colored boxes on the left indicate the six major clusters and correspond to the colors shown in (b), (c), and (d). In (b) and (c), the positions that correspond to each of the six clusters are shown as spheres in the corresponding color on the CYP2C9 crystal structure (PDB: 1r9o). Alternate viewpoint shown in (c). In (d), relative solvent accessibility of each cluster shown as a box plot.

Conversely, positions comprising Clusters 4 and 6 were tolerant to substitution and were located on the surface of the protein, though Cluster 6 was more sensitive to charged and proline amino acid substitutions. Cluster 5 contained many positions in the transmembrane domain, not shown in the crystal structure. Cluster 5 also included part of the F-G loop, which defines a portion of the CYP2C9 substrate access channel and interacts with the membrane^50^. Substitutions in the transmembrane domain (positions 1-20) had little effect on activity, but had a larger effect on abundance. The largest effects in the transmembrane domain were from charged substitutions, which rarely occur in transmembrane domains. CYP2C9 is co-translationally inserted into the ER and the N-terminal transmembrane domain is involved in ER retention^51^, so substitutions in the transmembrane domain that impacted abundance may have caused mislocalization.

### Predicting the clinical impact of human CYP2C9 variants

Genetic variation in CYP2C9 can drive variable drug response, but most CYP2C9 variants documented in humans so far have unknown functional consequences. The best-studied set of CYP2C9 variants are the 70 star alleles in PharmVar, some of which have been functionally characterized. The Clinical Pharmacogenetics Implementation Consortium (CPIC) reviews functional evidence and has made clinical recommendations for 35 of the 63 CYP2C9 single amino acid star alleles^16, 17^. We compared CPIC recommendations to our activity classes and found that CPIC allele function classes were largely concordant with our CYP2C9 activity classes (Figure 6). The few cases where our activity classes did not match CPIC classes were generally due to alleles with limited or inadequate functional evidence, as determined by CPIC (Supplementary Table 7).

**Figure 6.**
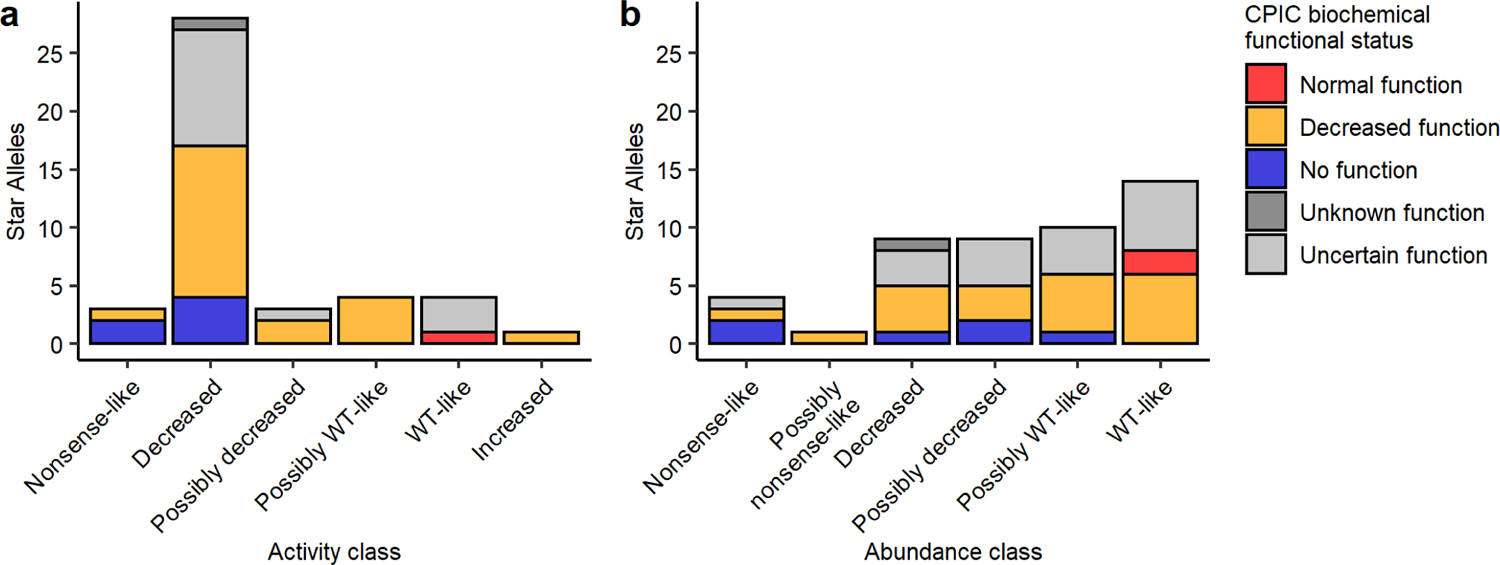
Comparison of activity and abundance scores with clinical pharmacogenomic recommendations. Stacked bar plot of number of CYP2C9 star alleles vs a) activity or b) abundance class, colored by clinical pharmacogenomic recommendation (CPIC biochemical functional class status). CPIC classes are taken from NSAID clinical functional status recommendations^16^.

We additionally curated 629 synonymous, missense, and nonsense CYP2C9 variants from the gnomAD database^20^, 559 of these with at least one functional score from our datasets. Most of these variants lack functional annotations, as only 27 of them are star alleles with an associated CPIC functional recommendation. All 8 nonsense variants had very low activity and/or abundance scores, all 119 synonymous variants had high activity and/or abundance scores, and missense variants spanned the range of activity and abundance scores (Supplementary Figure 9). Of the 466 total missense variants in gnomAD, 340 had an activity score (319 of these lack a CPIC functional recommendation), and a majority of these had significantly decreased activity. 58.8% of missense variants (200 variants) had “decreased” or “possibly decreased” activity, and 9.7% (33 variants) had “nonsense-like” or “possibly nonsense-like” activity (Figure 7). 168 of the missense variants were singletons in gnomAD, and these had the same activity score pattern as the other missense variants where 60.1% (101 variants) had “decreased” or “possibly decreased” activity, and 11.9% (20 variants) had “nonsense-like” or “possibly nonsense-like” activity. Finally, we compared our scores to several widely used computational predictors and found only moderate correlation between predicted functional status and experimentally derived activity scores (mean absolute Pearson’s *r*=0.494, Supplementary Figure 10). The fact that many human CYP2C9 variants have significantly decreased function is striking, highlighting that a large proportion of all possible CYP2C9 variants have the potential to impact the metabolism of warfarin and other drugs.

**Figure 7.**
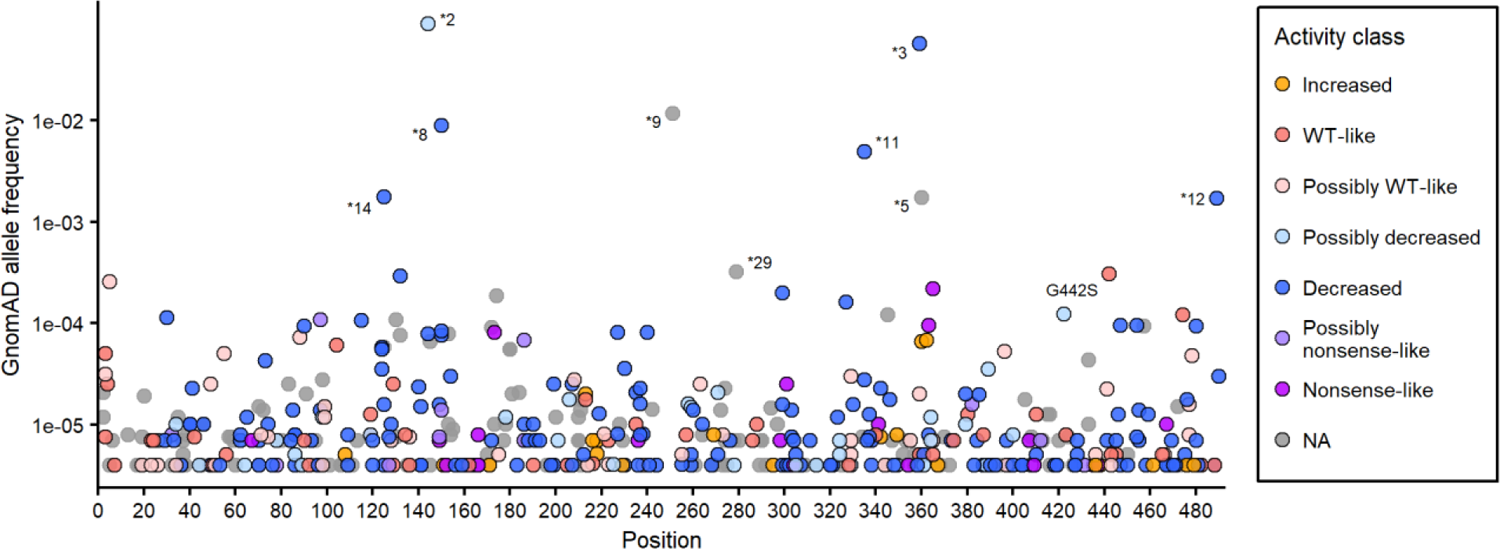
Classification of human CYP2C9 variants using activity data. Frequency and protein position of CYP2C9 missense variants in human population database gnomAD, colored by Click-seq activity class. Allele frequencies were calculated from combined v2 and v3 gnomAD allele frequencies. Variants at population frequency greater than 3×10^-4^ are labeled by star allele (if applicable) or amino acid change. Human variants lacking an activity class are shown in grey.

## DISCUSSION

CYP2C9 is a well-studied metabolic enzyme, and many small-scale functional characterizations of CYP2C9 have been performed^52–54^, with the largest of these comprising 109 CYP2C9 variants profiled for abundance using a VAMP-seq style assay^55^. Abundance scores from this study correlate well with our abundance scores (Pearson’s *r*=0.74). Collectively, previous studies of CYP2C9 variant function have tested only a small fraction of the possible single mutations, focusing only on already observed alleles. Therefore, we developed a high-throughput yeast activity assay, applied VAMP-seq, and generated activity and abundance scores for a combined total of 8,091 missense variants, or 87% of the possible missense variants in CYP2C9. Our results were highly reproducible across biological replicates and validated well when tested against individual variants and clinical substrates of CYP2C9. Additionally, our activity scores were concordant with CYP2C9 star allele functional status recommendations from CPIC^16^. In addition to CYP2C9 alleles of known function, we generated functional scores for over 300 CYP2C9 missense alleles present in gnomAD that currently lack functional annotation.

Our functional scores reflected known structural features of CYPs, including heme-binding residues and the highly conserved core regions of the protein. In general, residues involved in heme coordination and binding were crucial for activity, but were less important to protein abundance, indicating that heme insertion, a process which is not fully understood, is not necessarily stabilizing^56^. Somewhat surprisingly, residues in the active site were fairly tolerant to substitution and largely did not result in large decreases in activity. Instead, we found that substitutions in the hydrophobic core of the protein comprising helices D, E, I, J, K, and L were crucial for protein abundance and activity, and substitutions in these regions likely most affect protein stability.

We observed a strong correlation between activity and abundance scores, which is in contrast to other deep mutational scans. Paired activity and abundance data has been collected for VKOR and NUDT15^26, 27^, and for these proteins activity and abundance scores were much less well correlated (VKOR: Pearson’s *r* = 0.261, Spearman’s *ρ* = 0.25; NUDT15: Pearson’s *r* = 0.384, Spearman’s *ρ* = 0.34). The strong correlation of CYP2C9 activity and abundance scores is partially due to the design of the Click-seq vector, which does not include an expression control. Overall, we estimated that protein abundance could explain about 50% of the variation in CYP2C9 variant activity.

The variant functional datasets we generated are a resource for improving genotype-based dosing and also for improving our understanding of CYP biology. However, there are limitations to keep in mind. First, in both systems we expressed CYP2C9 as a cDNA using an inducible promoter, so our assays did not capture splicing defects or transcriptional regulation. CYP2C9 is constitutively expressed in the liver, but is also inducible via a number of substrates^9^. We also cannot discern the impact of protein interactions such as with CYP accessory proteins cytochrome P450 reductase (CPR) and cytochrome b5, or with other CYP enzymes. Additionally, due to our flow cytometry binning strategy, we suspect that Click-seq labeling was saturated at increased activity levels, so we would likely need to re-sort our library with a modified binning strategy to detect variants with significantly increased activity. An example of this is the G442S variant, which had a “WT-like” activity score but showed 130% and 180% wild type activity in individual tests (Figure 2), although this could also be explained by a substrate-dependent effect. Despite our binning strategy, we observed 240 variants with increased function, and 20 of these were also present in gnomAD. One increased activity variant, I434F (*59), was also present in the PharmVar database, warranting further investigation, especially since there are no documented CYP2C9 increased activity alleles^14^, so the clinical impact of an increased activity variant is unknown.

Finally, Click-seq measures CYP2C9 activity using a single substrate, so we cannot say how many variants may exhibit substrate-dependent effects, for which there is some evidence^57^. As an example, Arg108 has been shown to be critical for binding negatively charged substrates^35^ and for binding flurbiprofen in particular^42^, but substitutions at this position had little effect on activity as measured by Click-seq or abundance. This is likely because the TAHA probe is an amide and not acidic, so is able to bind regardless of a strong electrostatic interaction with Arg108. Despite this, our activity scores correlate well with S-warfarin and phenytoin activity overall, indicating that they are generally informative of a larger set of substrates. In the future, we plan to re-test our library with a range of activity-based probes to identify variants that result in substrate-dependent changes in function.

We also anticipate that Click-seq can be leveraged to examine other CYPs important to human drug metabolism such as CYP2D6 and CYP2C19, both of which have also been successfully expressed in yeast previously and for which activity-based click probes have been designed^29^. Expanding the repertoire of CYP deep mutational scans will allow us to investigate differences between CYP isoforms that are key to human drug metabolism and pharmacogenomics. Moreover, activity-based click probes are available for a variety of enzyme activities, so Click-seq is likely to be useful beyond CYPs.

In addition to revealing details of how CYP2C9 sequence relates to its structure and function, we hope that the variant functional data presented here will be useful for informing drug dosing. In particular, we hope that the data will empower CPIC to provide functional classifications for additional variants, perhaps preemptively. Such preemptive classification could be extremely powerful, as clinical genotyping efforts are increasing, guaranteeing we will continue to find new variants that may have consequences for drug dosing.

## METHODS

### General Reagents

Unless otherwise noted, all chemicals were obtained from Sigma Aldrich Chemical Co. (St. Louis, MO) and all enzymes were obtained from New England Biolabs. Tienilic acid, 6-hydroxywarfarin-d_5_, 7-hydroxywarfarin-d_5_ and 4-hydroxyphenytoin-d_5_ were synthesized according to published protocols^58^. Hex-5-yn-1-amine was purchased from GFS Chemicals (Powell, OH).

### Strains, plasmids, and oligonucleotides

All yeast strains are listed in Supplementary Table 1. All plasmids and oligonucleotides are listed in Supplementary Table 4

### Growth media and culturing techniques

*E. coli* were cultured at 37°C in Luria broth. Yeast were cultured at 30°C. Yeast culture media was prepared according to the following recipes. YP: 1% yeast extract, 2% peptone. YPD: 1% yeast extract, 2% peptone, 2% (w/v) glucose. C-ura: Yeast nitrogen base without amino acids and ammonium sulfate: 0.17%, ammonium sulfate: 0.5%, dropout mix lacking uracil: 0.2%, 2% (w/v) glucose. C-trp: Yeast nitrogen base without amino acids and ammonium sulfate: 0.17%, ammonium sulfate: 0.5%, dropout mix lacking tryptophan: 0.2%, 2% (w/v) glucose. Unless otherwise specified, all yeast transformations were performed using the LiAc/SS carrier DNA/PEG method^59^.

Yeast cells carrying *CYP2C9* wild type or variant plasmid were induced as follows: a single colony was inoculated into 5 mL YPD media supplemented with 200 μg/mL G418 and grown overnight with rotation. This culture was diluted 1:50 into fresh YP media containing 2% (w/v) raffinose and supplemented with 200 μg/mL G418 and grown for at least two cell doublings. Cultures were then inoculated to OD 0.0125 into fresh YP media containing 2% (w/v) galactose and 200 μg/mL G418 and collected after 7 doublings the following day.

All cell culture reagents were purchased from ThermoFisher Scientific unless otherwise noted. HEK 293T cells (ATCC CRL-3216) and derivatives thereof were cultured in Dulbecco’s modified Eagle’s medium supplemented with 10% fetal bovine serum, 100 U/mL penicillin, and 0.1 mg/mL streptomycin. Cells were induced with 2.5 µg/mL doxycycline. Cells were passaged by detachment with trypsin-EDTA 0.25%, and cells were prepared for sorting by detachment with versene. All cell lines tested negative for mycoplasma.

### Yeast strain engineering

A previously generated S288C derivative strain YMD3289 (*MATɑ HAP1+ ura3Δ0 leu2Δ1 his3Δ1 trp1Δ63*)^60^ was engineered to have improved human P450 activity by increasing protein expression and by expressing human CYP accessory proteins cytochrome P450 reductase (CPR) and cytochrome b5. First, the vacuolar protease genes *PEP4* and *PRB1* were sequentially knocked out to improve protein expression using the pop-in pop-out method^61^ using the vector pRS406^62^ with flanking sequences cloned in, resulting in the strain YMD4253. Next, *S. cerevisiae* codon-optimized *POR* sequence (human CPR) (Uniprot: P16435) was synthesized (Integrated DNA Technologies) with a C-terminal FLAG tag (sequence: DYKDDDDK) and cloned into a low-copy p416*GAL1* vector^63^, resulting in the plasmid p416*GAL1-hCPR*-*FLAG*. The auxotrophic marker *TRP1* was amplified from pRS414^62^ and cloned into this vector, and the fragment containing both *GAL1pr*::*hCPR*-*FLAG* and *TRP1* was amplified, digested with DpnI (NEB R0176), and used to transform the yeast strain YMD4253, resulting in strain YMD4254. Transformants were selected for growth on synthetic media lacking tryptophan (C-trp). Finally, *S. cerevisiae* codon-optimized cytochrome *b5* sequence (Uniprot: P00167) was synthesized (Integrated DNA Technologies) with an N-terminal MYC tag (sequence: EQKLISEEDL) and cloned into a low-copy p416*GPD* vector^64^. A portion of the vector containing both *GPDpr::MYC-hb5* and *URA3* was amplified via PCR, digested with DpnI, and used to transform yeast strain YMD4254, resulting in strain YMD4255. Transformants were selected for growth on synthetic media lacking uracil (C-ura). The fully humanized strain YMD4255 was backcrossed twice with YMD4252, resulting in the strain YMD4256 with genotype *MATa ura3Δ0::GPDpr::MYC-hb5::URA3 leu2Δ1 his3Δ1 trp1Δ63 HAP1+ pep4Δ0 prb1Δ0 ho::GAL1pr::hCPR-FLAG::TRP1* (strain details in Supplementary Table 1).

The low-copy p41K*GAL1* vector was constructed from the p416*GAL1* vector^63^ and the pUG6 vector^65^ using Gibson assembly^66^ to clone KanMX into p416*GAL1*. *S. cerevisiae* codon-optimized *CYP2C9* sequence (Uniprot: P11712) was synthesized (Integrated DNA Technologies) with a C-terminal HA tag (sequence: YPYDVPDYA) and cloned into p41K*GAL1* using Gibson assembly. Yeast strain YMD4256 was transformed with p41K*GAL1*-*hCYP2C9*-*HA* using the standard LiAc protocol referenced above and transformants were selected on YPD media supplemented with 200 µg/mL G418 to maintain the plasmid.

### *CYP2C9* yeast codon-optimized variant library construction in *S. cerevisiae*

The yeast *CYP2C9* variant library was generated using an inverse PCR-based site-directed saturation mutagenesis approach^67^. Saturation mutagenesis primers were designed for each codon in *CYP2C9* from positions 2 to 490 such that the forward primer contained an NNK at the 5’ end of the sequence. Primers were ordered resuspended from IDT. Forward and reverse primers for each codon position were mixed and inverse PCR was performed using 2.5 µM primers, 5% DMSO, 125 pg of *CYP2C9* template sequence, and KAPA Hifi Hotstart 2X ReadyMix (KAPA Biosytems KK2601). To generate the template *CYP2C9* vector, the *S. cerevisiae* codon-optimized *CYP2C9* sequence from p41K*GAL1-hCYP2C9*-*HA* was cloned into pHSG298 (Clontech) using restriction sites SalI and XbaI.

After inverse PCR was performed for every position, amplified variant constructs were verified by gel electrophoresis, quantified by Qubit fluorometry (Life Technologies), and pooled at equimolar ratios. Variant fragments were then treated with T4 polynucleotide kinase (NEB M0201) and ligated with T4 DNA ligase (NEB M0202) before transforming electrocompetent *E. coli* cells (NEB C2989K) with the ligated products, selecting for kanamycin resistance, and midiprepping (Qiagen). Next, the library was transferred from the pHSG298 vector used for saturation mutagenesis to the p41K*GAL1* yeast expression vector using an antibiotic switching strategy. The library was subcloned back into the low-copy p41K*GAL1* vector using restriction sites SpeI and SalI and ligated products were used to transform electrocompetent *E. coli* cells (NEB C2989K), selecting for growth on LB + ampicillin, and midiprepped (Qiagen).

To barcode the library, plasmid harvested from midiprep was digested with SalI at 37°C for 1 hour, and heat inactivated at 65°C for 20 minutes. Barcode oligos with 18 bp random sequences were ordered from IDT, resuspended at 100 µM, and then annealed by combining 1 µL each of primer with 4 µL CutSmart Buffer and 34 µL ddH2O and running at 98°C for 3 minutes followed by ramping down to 25°C at −0.1°C/second. After annealing, 0.8 µL of Klenow polymerase (exonuclease negative, NEB) and 1.35 µL of 1 mM dNTPS was then combined with the 40 µL of product to fill in the barcode oligo (cycling conditions: 25°C for 15 minutes, 70°C for 20 minutes, ramp down to 37°C at −0.1°C/s). Digested vector and barcode oligo were then ligated overnight at 16°C. The barcoded library was used to transform electrocompetent *E. coli* cells (NEB C2989K) and midiprepped (Qiagen). The size of the barcoded library was estimated using colony counts to be 280,000. To reduce library size, the barcoded library was again used to transform electrocompetent *E. coli* cells (NEB C2989K), bottlenecked, and midiprepped (Qiagen). The size of the barcoded library was estimated using colony counts to be 42,000.

To determine more accurate library barcode counts, 2 PCR replicates each using 1.5 µg of plasmid extracted library were amplified using custom barseq primers CJA120/CJA138 using KAPA2G Robust HotStart ReadyMix (Sigma 2GRHSRMKB) with the following conditions: 95°C for 3 m, 5 cycles of 95°C for 15 s, 60°C for 15 s, 72°C for 15 s, and 72°C for 1 m, then purified using AMPure XP beads (Beckman Coulter A63880) at 1:1 ratio (beads:DNA). The purified products were amplified using primers CJA135 and JS486 or JS487 using KAPA2G Robust HotStart ReadyMix with the following PCR conditions: 95°C for 3 m, 10 cycles of 95°C for 15 s, 65°C for 15 s, 72°C for 15 s, and 72°C for 1 m, and then gel extracted using the QIAquick Gel Extraction Kit (Qiagen) and quantified by Qubit fluorometry (Life Technologies). PCR replicates were pooled at equimolar ratios and deep sequenced on an Illumina NextSeq500 to determine the number of barcodes present. Briefly, forward and reverse reads were merged with Pear^68^, barcodes were counted with Enrich2^69^, and barcodes with less than 10 reads were removed, resulting in a total of ∼160,000 unique barcodes in the *CYP2C9* library, for an average of 17x coverage.

The barcoded *CYP2C9* library was used to transform the humanized yeast strain YMD4256, using the standard high-efficiency LiAc procedure mentioned above. Four independent transformations were pooled to generate a library stock of OD_600_ 5.7, equivalent to an average of 11x coverage (independent transformants) for each of the 160,000 independent barcoded variants. The latter estimate assumes that each yeast cell harbors one *CYP2C9* variant and that all growth rates are similar. Library stocks were stored at −80°C in 25% (v/v) glycerol.

### *CYP2C9* human library construction in HEK293T cells

*CYP2C9* sequence (Uniprot: P11712) codon-optimized for human expression was synthesized (Integrated DNA Technologies) and cloned into the vector pHSG298. As with the yeast activity library, saturation mutagenesis primers were designed for each codon in *CYP2C9* from positions 2 to 490 and ordered resuspended from IDT. Forward and reverse primers for each position were mixed at 2.5 µM and used in a PCR reaction with 125 pg of template, 5% DMSO, and 5 µL of KAPA Hifi Hotstart 2X ReadyMix. PCR products were visualized on a 0.7% agarose gel to confirm amplification of the correct product. PCR products were then quantified using the Quant-iT PicoGreen dsDNA Assay kit (Invitrogen) using DNA control curves done in triplicate and pooled at equimolar ratios. Pooled PCR products were cleaned and concentrated using Zymogen Clean and Concentrate kit and then gel extracted. The pooled library was phosphorylated with T4 PNK (NEB), incubated at 37°C for 30 minutes, and heat inactivated at 65°C for 20 minutes. 8.5 µL of this phosphorylated product was combined with 1 µL of 10X T4 ligase buffer (NEB) and 0.5 µL of T4 DNA ligase (NEB) to make a 10 µL overnight ligation reaction. This reaction was incubated at 16°C overnight.

The overnight ligation was then cleaned and concentrated (Zymogen) and eluted in 6 µL of ddH2O. 1 µL of this ligation was then used to transform high efficiency *E. coli* (NEB C3020K) using electroporation (settings: 2 kV). Each reaction contained 1 µL of ligation (or ligation control or pUC19 10 pg/µL) and 25 µL of *E. coli*. 975 µL of pre-warmed SOC media was added to each cuvette after electroporation, transferred to a culture tube, and recovered at 37°C, shaking for 1 hour. At 1 hour, 1 and 10 µL samples from all cultures were taken and plated on appropriate media (LB + kanamycin for ligation and ligation control; LB + ampicillin for pUC19), the remaining 989 µL was used to inoculate a 50 mL culture (LB + kanamycin). Plates and 50 mL culture were incubated at 37°C overnight (shaking for 50 mL culture). Colonies on plates were then counted, and counts were used to calculate how many unique molecules were transformed to gauge coverage of the library. 50 mL culture was spun down and midiprepped.

To transfer the library from pHSG298 to the recombination vector (attB-CYP2C9-EGFP-IRES-mCherry), the pHSG298 library and recombination vector were digested with MluI and SphI for 1 hour at 65°C. The library and cut vector were then gel extracted. The library was then ligated with the cut vector at 5:1 using NEB T4 ligase, overnight at 16°C. The ligation was heat inactivated the next morning, and cleaned and concentrated with the Zymo kit. Another high efficiency transformation was performed the same as described above, except this ligation was plated on LB + ampicillin (antibiotic switching strategy). Plates and 50 mL culture were incubated at 37°C overnight (shaking for 50 mL culture). Colonies on plates were then counted, and counts were used to calculate how many unique molecules were transformed to gauge coverage of the library. 50 mL culture was spun down and midiprepped.

The library was barcoded using the same method as the yeast activity library but using the AgeI site for barcode insertion. The overnight barcode ligation was cleaned and concentrated and eluted in 6 µL of ddH2O. 1 µL of this ligation was then transformed into high efficiency *E. coli* using electroporation at 2 kV. Each reaction contained 1 µL of ligation (or ligation control or pUC19 10 pg/µL) and 25 µL of *E. coli*. 975 µL of pre-warmed SOC media was added to each cuvette after electroporation, transferred to a culture tube, and recovered at 37°C, shaking for 1 hour. At 1 hour, 1 and 10 µL samples from water and pUC19 cultures were taken and plated on LB supplemented with ampicillin. For ligation and ligation control, four flasks were prepared with 50 mL of LB and ampicillin, and then 500 µL, 250 µL, 125 µL, and 62.5 µL was sampled from the 1 mL of recovery and transferred into a corresponding flask. From those flasks, 1 µL, 10 µL, and 100 µL, were sampled and plated onto LB ampicillin plates. Plates and 50 mL culture were incubated at 37°C overnight. Colonies on plates were then counted, and counts were used to calculate how many unique molecules were transformed to gauge the number of barcodes. Flask with the target number of barcodes was then spun down and midiprepped.

### PacBio sequencing of *CYP2C9* libraries for barcode-variant mapping

PacBio libraries were generated using the SMRTbell Express Template Prep Kit 2.0 (Pacific Biosciences) according to manufacturer’s directions with the following modifications. Barcoded variant sequences were excised using SpeI-HF and PspXI (activity library) or NheI and SmaI (abundance library) restriction enzymes and purified using AMPure PB beads (Pacific Biosciences 100-265-900) at 1:1 ratio (beads:DNA). Following end-repair and blunt end adaptor ligation, according to manufacturer’s instructions, PacBio libraries were subject to 2 additional rounds of restriction digestion to remove any backbone plasmid contamination present in the library. Finally, libraries were cleaned in 3 consecutive rounds of AMPure PB beads (Pacific Biosciences 100-265-900) at 0.6:1 ratio (beads:DNA). The purity and size of Pacbio libraries were confirmed by Tapestation (Agilent) and Bioanalyzer 2100 (Agilent) before proceeding with the sequencing run. Samples were submitted to University of Washington PacBio Sequencing Services and sequenced on two SMRT cells per library in a Sequel run. The yeast activity library was sequenced using two replicate library preparations from the same miniprep, while the human abundance library was sequenced from two replicate samplings of the same *E. coli* ligation transformation.

Long reads were filtered for at least 10 passes and analyzed using a custom analysis pipeline to identify and link gene and barcode regions (https://github.com/shendurelab/AssemblyByPacBio). The activity library contained 66,958 unique nucleotide variants (22,421 of these full-length, aka without indels), tagged by 105,372 unique barcodes, while the abundance library contained 37,758 unique nucleotide variants (22,669 of these full-length), tagged by 78,740 unique barcodes (Supplementary Table 3).

### Tienilic Acid Hexynyl Amide (TAHA) synthesis (activity-based probe)

Tienilic Acid (50 mg, 0.15 mmol), EDC (36 mg, 0.18 mmol) and 1-hydroxybenzotriazole hydrate (25 mg, 0.18 mmol), stirring under a nitrogen atmosphere at room temperature, were dissolved in 1 mL of anhydrous acetonitrile and 0.5 mL of anhydrous N,N-dimethylformamide. N-Methylmorpholine (56 μL, 0.45 mmol) was added and the reaction was stirred 15 minutes prior to the addition of hex-5-yn-1-amine (27 μL, 0.18 mmol). The reaction was then stirred another 4 hours after which it was diluted with ethyl acetate and successively washed with 10 % saturated sodium bicarbonate, water, and brine. The organic phase was dried over MgSO_4_ and solvent was evaporated. The final product was purified by flash chromatography, using a hexane/ethyl acetate gradient, and was obtained as a clear oil (52 mg, 84 % yield).

^1^H NMR spectra was recorded at 25°C in deuterated methanol (CD_3_OD) on a 500 MHz Agilent DD2 (Santa Clara, CA) spectrometer, (500 MHz, CD_3_OD): δ 8.00 (d, *J* = 4.40 Hz, 1H), 7.48 (d, *J* = 4.40 Hz, 1H), 7.46 (d, *J* = 8.79 Hz, 1H), 7.21 (t, *J* = 4.40 Hz, 1H), 7.14 (d, *J* = 8.79 Hz, 1H), 4.74 (s, 2H), 3.35 (t, *J* = 6.83 Hz, 2H), 2.26-2.21 (m, 3H), 1.70 (quin, *J* = 6.83 Hz, 2H), 1.56 (quin, *J* = 6.83, 2H). ^1^H-decoupled ^13^C NMR (^13^C{^1^H}) spectra was recorded at 25°C in acetone-d_6_ (C_3_D_6_O) on a 500 MHz Bruker Avance DRX-500 (Billerica, MA) spectrometer, equipped with a Bruker triple resonance TXO probehead. Chemical shifts are reported below relative to the solvent peaks in C_3_D_6_O at 206.7 and 29.9 ppm. ^13^C{^1^H} NMR (125 MHz, C_3_D_6_O) δ 186.0, 167.3, 156.9, 144.5, 137.0, 136.9, 133.9, 131.3, 129.6, 128.6, 123.4, 112.8, 84.7, 70.1, 69.4, 39.0, 29.5, 26.6, 18.4.

High resolution mass spectrometry (HRMS) was determined via UPLC-MS on a Waters Acquity UPLC (Milford, MA) coupled to an AB Sciex TripleTOF 5600 mass spectrometer (Framingham, MA). Data analysis was performed with AB Sciex Analyst TF 1.7.1. HRMS (ESI+) m/z [M + H] calculated (C_19_H_18_Cl_2_NO_3_S) 410.0379, observed 410.0374, δ ppm 1.22. All NMR and mass spectra have been provided in the Supplementary Information section (Supplementary Figure 12 and Supplementary Figure 13).

### FACS-based deep mutational scan of activity library (Click-seq)

#### CYP2C9 activity assay with CYP2C9-specific probe (activity-based protein profiling)

CYP2C9 enzymatic activity was probed using a flow-cytometry based method with a click chemistry compatible probe TAHA-ABP (synthesis described above) that has specificity for CYP2C9 activity with minimum reactivity towards other yeast proteins. Yeast cultures were grown as described above to induce CYP expression, and for each sample, 1 OD of overnight yeast culture was collected via centrifugation at 4000 rpm for 2 min, washed with 0.5 mL of PBS by resuspension and centrifugation at 4000 rpm for 2 min, and resuspended in 100 µL PBS:0.1% saponin (w/v). Each sample was pre-incubated with 2 mM NADPH (Sigma N1630) at 37°C for 20 mins. All samples except a ‘No probe’ control were treated with 10 µM TAHA-ABP and incubated with rotation at 37°C for 20 hrs to form activity-dependent CYP2C9-probe adducts. Samples were collected via centrifugation at 4000 rpm for 2 min and washed three times with 0.5 mL PBS as above. Samples were resuspended in 100 µL PBS:0.1% saponin (w/v) and incubated at room temperature for 20 mins. 100 µL 2x copper-catalyzed azide-alkyne cycloaddition (CuAAC) reaction buffer was added to cells to append a fluorophore reporter (2x concentrations: 10 µM CF488A picolyl azide (Biotium #92187), 2 mM CuSO_4_ (Sigma C8027), 4 mM THPTA (Sigma 762342), 6 mM ascorbic acid (Sigma A7631) in PBS) and vortexed vigorously to mix. Samples were incubated in the dark at room temperature for 30 minutes and collected by centrifugation as above. Cells were washed five times in 0.5 mL PBS, resuspended in 1 mL PBS, and stored at 4°C up to 1 day.

### CYP2C9 library labeling and FACS

To label and sort the CYP2C9 yeast library, isogenic humanized yeast strains expressing control CYP2C9 variants (wild type, R144C, I359L, and C435H) were induced in galactose as described above. The barcoded *CYP2C9* variant library was thawed at room temperature and ∼8 OD of library was inoculated into 25 mL YPD media supplemented with 200 µg/mL G418 and grown overnight at 150 rpm. The rest of the induction was performed as described above, with 5x culture volumes and shaking instead of rotation. For each control variant, one sample was collected (1 OD), and for the library, 4 samples (1 OD each) were collected. A “no probe” sample was included as a control. All samples were labeled using the activity assay described above with CYP2C9-specific activity-based probe TAHA-ABP.

Labeled cells were run on a BD AriaIII sorter (BD Bioscience, San Jose, CA) and a standard yeast singlet gate was used. For this population, data were collected on the AF488A channel (488 nm excitation; 530/30 nm detection filter), and gates were drawn to contain 10%, 10%, 20%, and 60% of events from the library sample, from most fluorescent (AF488A channel) to least fluorescent. Gates were sorted into 5 mL tubes, harvested by centrifugation and stored at −20°C before library preparation. Flow cytometry data were collected using FACSDiva version 8.0.1 (BD Biosciences). See Supplementary Table 2 for details of numbers of cells collected. Four biological replicates of the FACS-based deep mutational scan were performed.

#### CYP2C9 sorted activity library amplification and sequencing

For the yeast activity library, sorted samples were harvested by centrifugation and stored at −20°C. Plasmids were extracted from sorted cell pellets using the Zymoprep Yeast Plasmid Miniprep I kit (Zymo Research D2001). Each sorted sample was split into two for PCR replicates. For each sample, the barcode region was amplified and an 18bp unique molecular identifier (UMI) sequence was added using primers CJA120/CJA124 using KAPA2G Robust HotStart ReadyMix with the following conditions: 95°C for 3 m, 2 cycles of 95°C for 20 s, 60°C for 15 s, 72°C for 30 s, and 72°C for 1 m, then purified using AMPure XP beads (Beckman Coulter A63880) at 1:1 ratio (beads:DNA). Purified products were amplified using various forward (CJA135, CJA139, or CJA144) and reverse indexing primers (JS409-412,JS470-477) using KAPA2G Robust HotStart ReadyMix with 0.5x SYBR green (Roche #04707516001) on a miniOpticon (Bio-Rad) with the following PCR conditions: 95°C for 3 m, up to 30 cycles of 95°C for 20 s, 65°C for 15 s, 72°C for 30 s, and removed from the thermocycler when the relative fluorescence units (RFU) was between 0.5 and 1. These products were again purified using AMPure XP beads (Beckman Coulter A63880) at 1:1 ratio (beads:DNA), and were then gel extracted using the QIAquick Gel Extraction Kit (Qiagen) and quantified by Qubit fluorometry (Life Technologies). Samples were pooled at equimolar ratios and deep sequenced on an Illumina NextSeq500. Within each sort there was a good correlation of barcode frequencies from PCR replicates (mean Pearson’s *r* = 0.859, mean Spearman’s *ρ* = 0.694, Supplementary Figure 2).

### FACS-based deep mutational scan of abundance library

#### CYP2C9 abundance library transfection and FACS

HEK293T cells with a serine integrase landing pad integrated via lentivirus with a selectable inducible Caspase 9 cassette (HEK293T-LLP-iCasp9)^33^ were used for all human cell experiments, enabling expression of a single variant per cell. To recombine variants into HEK293T cells, cells were transfected in 10 cm plates, 3,500,000 cells per plate (4 plates per replicate). 7.1 µg of library plasmid was mixed with 0.48 µg of Bxb1 plasmid in 710 µL of OptiMEM. In a separate tube, 28.5 µL of Fugene was diluted in 685 µL of OptiMEM. The tubes were then combined and incubated at room temperature for 15 minutes. After incubation period, Fugene/DNA mixture was added to cells dropwise, and plates were placed in incubator at 37°C. A minimum of 48 hours after transfection, cells were induced with doxycycline at a final concentration of 2.5 µg/mL. 24 hours after induction with doxycycline, small molecule AP1903 was added to select from recombinant cells, which causes inducible Caspase 9 in unrecombined landing pads to dimerize and activate.

Recombined HEK293T cells were run on a BD AriaIII sorter. Cells were gated for live, recombined singlets. For this population, a ratio of eGFP/mCherry was calculated, and the histogram of this ratio was divided into four quartiles. Each quartile was sorted into a 5 mL tube. Sorted cells were grown out for 2-4 days post sorting to ensure enough DNA for sequencing. Three biological replicates of the FACS-based deep mutational scan were performed.

#### CYP2C9 sorted abundance library amplification and sequencing

For the abundance library, cells were collected, pelleted by centrifugation and stored at −20°C. Genomic DNA was prepared using a DNEasy kit, according to the manufacturer’s instructions (Qiagen), with the addition of a 30 min incubation at 37 °C with RNAse in the re-suspension step. Eight 50 μL first-round PCR reactions were each prepared with a final concentration of ∼50 ng/μL input genomic DNA, 1 × Q5 High-Fidelity Master Mix and 0.25 μM of the KAM499/VKORampR 1.1 primers. The reaction conditions were 98°C for 30 s, 98°C for 10 s, 65°C for 20 s, 72°C for 60 s, repeat 5 times, 72°C for 2 min, 4°C hold. Eight 50 μL reactions were combined, bound to AMPure XP (Beckman Coulter), cleaned and eluted with 21 μL water. Forty percent of the eluted volume was mixed with Q5 High-Fidelity Master Mix; VKOR_indexF_1.1 and one of the indexed reverse primers, JS385 through JS388, were added at 0.25 μM each. These reactions were run with Sybr Green I on a BioRad MiniOpticon; reactions were denatured for 3 minutes at 95°C and cycled 20 times at 95°C for 15 s, 60°C for 15 s, 72°C for 15 s with a final 3 min extension at 72°C. The indexed amplicons were mixed based in relative fluorescence units and run on a 1% agarose gel with Sybr Safe and gel extracted using a freeze and squeeze column (Bio-Rad). The product was quantified using KAPA Library Quant kit (KAPA Biosystems).

### Library sequence analysis

#### Activity library sequence analysis

For the activity library, barcode and UMI sequences were trimmed and filtered for minimum base quality Q20 using FASTX-toolkit (http://hannonlab.cshl.edu/fastx_toolkit/). Barcodes were collapsed according to UMIs by pasting the UMI sequence after the barcode sequence for each read, then identifying unique combinations of barcode-UMI (sort | uniq-c). The barcode from each unique barcode-UMI pair was used to generate a FASTQ files that was then input into Enrich2^69^ to count variants. Barcodes assigned to variants containing insertion, deletions, or multiple amino-acid alterations were removed from the analysis, and barcode counts were collapsed into variant counts. Variants were kept if they had a total (across bin) frequency greater than 1e-5 in each replicate (see Supplementary Figure 11). For each replicate, a weighted average of variant frequency across bins was used to determine activity score. To determine optimal bin weights, a linear regression on activity score (pool score) versus individual variant TAHA labeling was performed with 14 variants. Bin weights were varied to determine the best fit regression between pool score and individual score, resulting in the following bin weights: *w_1_ = 0.05* (bin1), *w_2_ = 0.2*, *w_3_ = 0.25*, *w_4_ = 1* (bin4), *R^2^ =* 0.986. Scores were normalized to the median synonymous weighted average (set to a score of 1), and the median nonsense weighted average of nonsense variants in the first 90% of the protein (score set to 0), and scores were averaged across replicates. Variants with less than two replicates were removed. Scores for missense variants range from −0.046 to 1.305 and have a bimodal distribution with peaks approximately matching the synonymous and nonsense distributions.

### Abundance library sequence analysis

For the abundance library, barcode sequences were trimmed and filtered for minimum base quality Q20 using FASTX-toolkit (http://hannonlab.cshl.edu/fastx_toolkit/). As with the activity library, barcodes were counted with Enrich2. Barcodes assigned to variants containing insertion, deletions, or multiple amino-acid alterations were removed from the analysis, and barcode counts were collapsed into variant counts. Variants were kept if they had a total (across bin) frequency greater than 1e-4 in each replicate (Supplementary Figure 11). Abundance scores were calculated as above, but with the following weights: *w_1_ = 0.25* (bin1), *w_2_ = 0.5*, *w_3_ = 0.75*, *w_4_ = 1* (bin4). Scores were normalized to the synonymous and nonsense distributions as above, but only normalizing to nonsense scores in the middle 80% of positions, excluding the first and last 10% of the protein. Variants with less than two replicates were removed. Scores for missense variants range from −0.29 to 1.59 and have a trimodal distribution with upper and lower peaks approximately matching the synonymous and nonsense distributions.

### Specific activity score calculations

To calculate specific activity score, activity and abundance scores were normalized such that the lowest and highest scores in the dataset were set to 0 and 1, respectively, and a ratio of normalized activity score to normalized abundance score was calculated (specific activity). Specific activity scores were only calculated for variants that had both activity and abundance scores.

### Classification of functional scores

Activity and abundance classes were determined as follows, based on a method modified from Matreyek et al. 2018 (Supplementary Figure 4). A synonymous score threshold was used to discriminate between ‘WT-like’ and ‘decreased’ scores. This threshold was set at the 5th percentile of synonymous scores (0.879 for activity score and 0.77 for abundance score). Variants were classified as ‘WT-like’ if their score and lower confidence interval were greater than the synonymous threshold, or ‘possibly WT-like’ if just their score was greater than the threshold. Variants were classified as ‘decreased’ if their score and upper confidence interval were less than the synonymous threshold, or ‘possibly decreased’ if just their score was less than the threshold. A nonsense score threshold was used to discriminate between ‘decreased’ and ‘nonsense-like’ scores, This threshold was the 95th percentile of nonsense scores (0.093 for activity score and 0.282 for abundance score). Variants were classified as ‘nonsense-like’ if their score and upper confidence interval were less than the nonsense threshold, or ‘possibly nonsense-like’ if just their score was less than the threshold. Finally, an upper synonymous threshold was used to discriminate between ‘WT-like’ and ‘increased’ scores, set at the 95th percentile of synonymous scores (1.102 for activity score and 1.212 for abundance score). Scores were classified as ‘increased’ if their score and lower confidence interval were greater than the upper synonymous threshold.

Analysis scripts are available at http://github.com/dunhamlab/CYP2C9. The Illumina and PacBio raw sequencing files and barcode–variant maps can be accessed at the NCBI Gene Expression Omnibus (GEO) repository under accession number GSE165412. The data presented in the manuscript are available as Supplementary Data files.

### Click-seq internal validation with individual CYP2C9 variants

14 individual *CYP2C9* variants were generated using an inverse PCR site-directed mutagenesis. Oligonucleotide pairs for each of the 14 variants are listed in Supplementary Table 4. With these, point mutations were generated using a KAPA HiFi DNA Polymerase (KAPA Biosytems KK2601) and 500 pg of *CYP2C9* template sequence p41K*GAL1-hCYP2C9*-*HA*. After performing inverse PCR for each variant, products were run on a 0.7% agarose gel, gel extracted using the QIAquick Gel Extraction Kit (Qiagen), treated with T4 polynucleotide kinase (NEB M0201) at 37°C for 30min, and ligated with T4 DNA ligase (NEB M0202) at 16°C overnight. Ligated products were used to transform chemically competent *E. coli* cells (NEB C2987 or Bioline BIO-85027). Bacterial clones were prepared for plasmid extraction using the QIAprep Spin Miniprep Kit (Qiagen) and variant sequences were confirmed with Sanger sequencing. Plasmids containing missense variants were individually transformed into YMD4256 using the 1-step transformation protocol^70^ and selection for growth in YPD supplemented with 200 µg/mL G418. Individual clones from each transformation were stored at −80°C.

Individual CYP2C9 yeast-expressed variants were grown and induced in galactose as described above, and 1 OD/mL of culture was collected for each variant. All samples were labeled using the CYP2C9 functional assay described above with CYP2C9-specific activity-based probe TAHA-ABP. Labeled cells were analyzed using a BD LSRII and a standard yeast singlet gate was used. For this population, data were collected on the FITC channel (488 nm excitation; 530/30 nm detection filter) for 20,000 events. Flow cytometry data were collected using FACSDiva version 8.0.1 (BD Biosciences) and analyzed using FlowJo version 10.7.1 (Ashland, OR). Fluorescence (FITC geometric mean of gated single cells) was normalized to background labeling (‘no probe’ control) and variant ABPP labeling relative to wild type was calculated. Three biological replicates of CYP2C9 individual variant validation were performed.

### VAMP-seq internal validation with individual CYP2C9 variants

12 of the 14 variants used for Click-seq validation that also had VAMP-seq abundance score were cloned using the IVA cloning^71^ site-directed mutagenesis method into the VAMP-seq recombination vector (attB-CYP2C9-EGFP-IRES-mCherry) using primers in Supplementary Table 4 (MAC379 through MAC403). Point mutations were generated using a KAPA HiFi DNA Polymerase (KAPA Biosytems KK2601) and 40 ng of *CYP2C9* template sequence attB-CYP2C9-EGFP-IRES-mCherry. After performing inverse PCR for each variant, products were digested with DpnI and were used to transform chemically competent *E. coli* cells (NEB C2987 or Bioline BIO-85027). Bacterial clones were prepped using a midiprep kit, validated by Sanger sequencing, and HEK293T-LLP-iCasp9 cells with landing pad were transfected with these preps. To recombine variants into HEK293T cells, cells were transfected in 6-well plates, 400,000 cells per well. 2.7 µg of library plasmid was mixed with 0.300 µg of Bxb1 plasmid in 125 µL of OptiMEM and 5 µL P3000 reagent. In a separate tube, 2.25 µL of Lipofectamine was diluted in 125 µL of OptiMEM. The tubes were then combined and incubated at room temperature for 15 minutes. After incubation period, Lipofectamine/DNA mixture was added to cells dropwise, and plates were placed in incubator at 37°C. A minimum of 48 hours after transfection, cells were induced with doxycycline at a final concentration of 2.5 µg/mL. 24 hours after induction with doxycycline, small molecule AP1903 was added to select for recombinant cells, which causes inducible Caspase 9 in unrecombined landing pads to dimerize and activate.

Recombined HEK293T cells were analyzed using a BD LSRII flow cytometer. Cells were gated for live, recombined singlets. For this population, a ratio of eGFP/mCherry was calculated, and the geometric mean of the histogram of this ratio was reported. Flow cytometry data were collected using FACSDiva version 8.0.1 (BD Biosciences) and analyzed using FlowJo version 10.7.1 (Ashland, OR). Two biological replicates of CYP2C9 individual variant validation were performed.

### Yeast microsomal preparations

A large-scale induction of yeast cells expressing *CYP2C9* wild type or variant plasmid was done as described above with a final culture volume of 0.5 L. After switching cells to galactose culture, cells were collected after 18-22 hrs. Cells were pelleted and stored at −80°C until ready for microsome preparations.

Yeast microsomes were prepared as described previously^60, 72^ with slight modifications. Harvested cells were thawed at room temperature for at least 10 minutes, washed with 25 mL of TEK buffer (50 mM Tris-HCl, pH 7.4, 1 mM EDTA, 0.1 M KCl), recovered at 3200 x g, resuspended in 30 mL of TEM buffer (50 mM Tris-HCl, pH 7.4, 1 mM EDTA, 70 mM 2-mercaptoethanol), and incubated at room temperature for 10 minutes. Cells were recovered by centrifugation (3200 x g) and resuspended in 1.5 mL of TMS buffer (1.5 M sorbitol; 20 mM Tris-MES, pH 6.3; 2 mM EDTA), and 20 mg of 20T Zymolyase (Amsbio) was added. Cells were incubated for ∼1 hour at 30°C with agitation until digested. Further steps were performed on ice. Spheroplasts were pelleted at 6732 x g and washed with 25 mL of TES-A buffer (50 mM Tris-HCl, pH 7.4, 1 mM EDTA, 1.5 M sorbitol), and the centrifugation step was repeated. Spheroplasts were resuspended in 10 mL of TES-B buffer (50 mM Tris-HCl, pH 7.4; 1 mM EDTA; 0.6 M sorbitol) and lysed using a Misonix S4000 by performing 4 x 15 second pulses at maximum amplitude (40-45 W). After 5 minutes on ice, lysed cells were centrifuged for 4 minutes at 1700 x g. The supernatant was then centrifuged at 110,000 x g for 70 minutes. The microsomal pellet was resuspended in 1 mL of TEG buffer (50 mM Tris-HCl pH 7.4, 1 mM EDTA, 20% (v/v) glycerol), homogenized, and frozen at −80°C.

### Warfarin metabolism validation assay

S-Warfarin (50 μM) was mixed together with yeast lysate (microsomes), prepared from CYP2C9 variant-expressing cells, at 5 mg/mL total protein in 100 mM KPi buffer, pH 7.4 (100 μL final incubation volume). After 3 minutes pre-incubation at 37°C in a water bath, NADPH was added to initiate (to 1 mM final concentration). Reactions were incubated for 20 minutes and were then quenched with the addition of 5 μL of ice-cold 70% HClO_4_. An internal standard solution, containing 5 ng each of 6-hydroxywarfarin-d_5_ and 7-hydroxywarfarin-d_5_, was added and the reaction products were vortexed and centrifuged to remove protein. Supernatants were analyzed by LC-MS/MS. Three technical replicates were carried out for each CYP2C9 variant lysate. Calibration curves were prepared by spiking variable amounts of unlabeled 6- and 7-hydroxywarfarins into 100 μL volumes of KPi buffer to generate standard mixtures with final concentrations ranging from 1 nM to 1 μM. These standard solutions were worked up and analyzed in an identical fashion to that described for the incubation samples.

### Phenytoin metabolism validation assay

Phenytoin (100 μM) was mixed together with yeast lysate (microsomes), prepared from CYP2C9 variant-expressing cells, at 5 mg/mL total protein in 100 mM KPi buffer, pH 7.4 (200 μL final incubation volume). After 3 minutes pre-incubation at 37°C in a water bath, NADPH stock was added (to 1 mM final concentration) to initiate the reactions. Reactions were incubated for 20 minutes and were then quenched with the addition of 20 μL of ice-cold 15% ZnSO_4_. 4-Hydroxyphenytoin-d_5_ (p-HPPH-d_5_, 10 ng) was added as the internal standard and the reactions were vortexed, then centrifuged to remove protein, and the supernatants were analyzed by LC-MS/MS. Again, three technical replicates were carried out for each CYP2C9 variant lysate. Calibration curves were prepared by spiking variable amounts of unlabeled 4-hydroxyphenytoin (p-HPPH) into 200 μL volumes of KPi buffer, generating standard mixtures with final concentrations ranging from 1 nM to 1 μM. These standard solutions were worked up and analyzed in an identical fashion to that described for the incubation samples.

### LC-MS/MS of Warfarin and Phenytoin Metabolites

LC-MS/MS analyses of warfarin and phenytoin metabolic reactions were conducted on a Waters Xevo TQ-S Tandem Quadrupole Mass Spectrometer (Waters Co., Milford, MA) coupled to an ACQUITY Ultra Performance LC™ (UPLC™) System with integral autoinjector (Waters). The Xevo was operated in ESI^+^-MS/MS (SRM) mode at a source temperature of 150°C and a desolvation temperature of 350°C. The following mass transitions were monitored in separate ion channels for the various oxidative warfarin metabolites/standards: *m/z* 325 > 179 (6- and 7-hydroxywarfarins-d_0_) and *m/z* 330 > 179 (6- and 7-hydroxywarfarins-d_5_); and phenytoin metabolite and standard: *m/z* 269 > 198 (p-HPPH-d_0_) and *m/z* 274 > 203 (p-HPPH-d_5_). Optimized cone voltages and collision energies were set to 25 V and 15 eV for all metabolites and standards of warfarin, while the cone voltage was set to 35 V with a collision energy of 15 eV for the phenytoin metabolite p-HPPH (both d_0_ and d_5_-labeled). Metabolic products from the warfarin incubations were separated on an Acquity BEH Phenyl, 1.7 μ, 2.1 x 150 mm UPLC column (Waters, Corp) using an isocratic gradient of 45% solvent A (0.1% aqueous formic acid) and 55% solvent B (methanol), with a constant flow rate of 0.35 mL/min. Phenytoin metabolites were separated using this same BEH Phenyl UPLC column with a solvent gradient of water (solvent A) and acetonitrile (solvent B), both of which contained 0.1% formic acid, running at a flow rate of 0.3 mL/min. Initially, solvent B was set to 28%, where it was maintained for 4.5 minutes, then increased linearly to 95% over 0.5 minutes where it was left for an additional 1.5 minutes. Metabolites were quantified through comparison of their peak area ratios (relative to either the 6- and 7-hydroxywarfarin-d_5_ or p-HPPH-d_5_ internal standard peak areas) to calibration curves using linear regression analysis. The limits of detection for all of the metabolites were below 5 fmol injected on column. Mass spectral data analyses for the Xevo TQ-S were performed on Windows XP-based Micromass MassLynxNT, v. 4.1, software (Waters).

### BOMCC fluorogenic assay with yeast microsomes

7-Benzyloxymethyloxy-3-cyanocoumarin (BOMCC) (50 μM) was mixed with 200 μM NADPH and yeast lysate at 50 μg total protein, prepared from CYP2C9 variant-expressing cells, in 50 mM KPi buffer, pH 8 (150 μL final incubation volume). Each sample was done in parallel with a no NADPH control. Three technical replicates were carried out for each CYP2C9 variant lysate. Sample fluorescence (excitation: 410 nm, emission: 460 nm, gain: 60) was recorded every 5 minutes on a BioTek Synergy H1 microplate reader at 37°C for 200 min with shaking. To determine relative activity, the fluorescence from each sample was normalized by subtracting the no NADPH control, and the slope of the normalized fluorescence signal during the linear range (5 mins to 50 mins) was calculated. Slopes were averaged across technical replicates and normalized by wild type average slope to determine relative BOMCC metabolism.

## ACKNOWLEDGEMENTS

We thank D. Prunkard of the UW Pathology Flow Cytometry Core Facility for assistance with cell sorting; K. Munson of the UW PacBio Sequencing Services for assistance with long-read sequencing; A. Wright, C. Whidbey, and E. Stoddard for reagents and helpful discussion; J. Stephany for assistance with amplicon design and sequencing analysis; D. Nickerson for valuable advice in analyzing the data, B. Dunn for useful comments in writing the manuscript; and all members of the Dunham lab for helpful feedback on figures. Research reported in this publication was supported by the National Institute of General Medical Sciences of the National Institutes of Health under award numbers R24GM115277 and R01GM132162 to DMF, MJD, and AER. CJA was supported by the National Human Genome Research Institute of the NIH under award T32 HG00035. The research of MJD was supported in part by a Faculty Scholar grant from the Howard Hughes Medical. MJD acknowledges prior support as a Senior Fellow in the Genetic Networks program at the Canadian Institute for Advanced Research.

## BIBLIOGRAPHY

1. Landrum, M. J. et al. ClinVar: Public archive of relationships among sequence variation and human phenotype. Nucleic Acids Res. 42, D980–D985 (2014).

2. Starita, L. M. et al. Variant Interpretation: Functional Assays to the Rescue. Am. J. Hum. Genet. 101, 315–325 (2017).

3. Sultana, J., Cutroneo, P. & Trifirò, G. Clinical and economic burden of adverse drug reactions. Journal of Pharmacology and Pharmacotherapeutics 4, S73–S77 (2013).

4. Lazarou, J., Pomeranz, B. H. & Corey, P. N. Incidence of adverse drug reactions in hospitalized patients: A meta-analysis of prospective studies. Journal of the American Medical Association 279, 1200–1205 (1998).

5. Budnitz, D. S. et al. National surveillance of emergency department visits for outpatient adverse drug events. J. Am. Med. Assoc. 296, 1858–1866 (2006).

6. Goulding, R., Dawes, D., Price, M., Wilkie, S. & Dawes, M. Genotype-guided drug prescribing: A systematic review and meta-analysis of randomized control trials. British Journal of Clinical Pharmacology 80, 868–877 (2015).

7. Zanger, U. M. & Schwab, M. Cytochrome P450 enzymes in drug metabolism: Regulation of gene expression, enzyme activities, and impact of genetic variation. Pharmacol. Ther. 138, 103–141 (2013).

8. Rettie, A. E. & Jones, J. P. Clinical and toxicological relevance of CYP2C9: Drug-drug interactions and pharmacogenetics. Annu. Rev. Pharmacol. Toxicol. 45, 477–494 (2005).

9. Daly, A. K., Rettie, A. E., Fowler, D. M. & Miners, J. O. Pharmacogenomics of CYP2C9: Functional and clinical considerations. Journal of Personalized Medicine 8, (2018).

10. Li, X. et al. Precision dosing of warfarin: open questions and strategies. Pharmacogenomics Journal 19, 219–229 (2019).

11. Steward, D. J. et al. Genetic association between sensitivity to warfarin and expression of CYP2C9*3. Pharmacogenetics 7, 361–367 (1997).

12. Pirmohamed, M., Kamali, F., Daly, A. K. & Wadelius, M. Oral anticoagulation: A critique of recent advances and controversies. Trends in Pharmacological Sciences 36, 153–163 (2015).

13. Sim, S. C. & Ingelman-Sundberg, M. The Human Cytochrome P450 (CYP) Allele Nomenclature website: a peer-reviewed database of CYP variants and their associated effects. Hum. Genomics 4, 278–281 (2010).

14. Gaedigk, A. et al. The Pharmacogene Variation (PharmVar) Consortium: Incorporation of the Human Cytochrome P450 (CYP) Allele Nomenclature Database. Clin. Pharmacol. Ther. 103, 399–401 (2018).

15. Relling, M. V. & Klein, T. E. CPIC: Clinical pharmacogenetics implementation consortium of the pharmacogenomics research network. Clinical Pharmacology and Therapeutics 89, 464–467 (2011).

16. Theken, K. N. et al. Clinical Pharmacogenetics Implementation Consortium Guideline (CPIC) for CYP2C9 and Nonsteroidal Anti-Inflammatory Drugs. Clin. Pharmacol. Ther. 108, 191–200 (2020).

17. Karnes, J. H. et al. Clinical Pharmacogenetics Implementation Consortium (CPIC) Guideline for CYP2C9 and HLA-B Genotypes and Phenytoin Dosing: 2020 Update. Clinical Pharmacology and Therapeutics (2020). doi:10.1002/cpt.2008

18. Zhou, Y., Ingelman-Sundberg, M. & Lauschke, V. M. Worldwide Distribution of Cytochrome P450 Alleles: A Meta-analysis of Population-scale Sequencing Projects. Clin. Pharmacol. Ther. 102, 688–700 (2017).

19. Gordon, A. S. et al. Quantifying rare, deleterious variation in 12 human cytochrome P450 drug-metabolism genes in a large-scale exome dataset. Hum. Mol. Genet. 23, 1957–1963 (2014).

20. Karczewski, K. J. et al. The mutational constraint spectrum quantified from variation in 141,456 humans. Nature 581, 434–443 (2020).

21. Fowler, D. M. & Fields, S. Deep mutational scanning: A new style of protein science. Nat. Methods 11, 801–807 (2014).

22. Romero, P. A., Tran, T. M. & Abate, A. R. Dissecting enzyme function with microfluidic-based deep mutational scanning. Proc. Natl. Acad. Sci. U. S. A. 112, 7159–7164 (2015).

23. Relling, M. V. et al. New Pharmacogenomics Research Network: An Open Community Catalyzing Research and Translation in Precision Medicine. Clin. Pharmacol. Ther. 102, 897–902 (2017).

24. Chiasson, M., Dunham, M. J., Rettie, A. E. & Fowler, D. M. Applying Multiplex Assays to Understand Variation in Pharmacogenes. Clin. Pharmacol. Ther. 106, 290–294 (2019).

25. Matreyek, K. A. et al. Multiplex assessment of protein variant abundance by massively parallel sequencing. Nat. Genet. 50, 874–882 (2018).

26. Suiter, C. C. et al. Massively parallel variant characterization identifies NUDT15 alleles associated with thiopurine toxicity. Proc. Natl. Acad. Sci. U. S. A. 117, 5394–5401 (2020).

27. Chiasson, M. A. et al. Multiplexed measurement of variant abundance and activity reveals VKOR topology, active site and human variant impact. Elife 9, 1–25 (2020).

28. Wright, A. T. & Cravatt, B. F. Chemical Proteomic Probes for Profiling Cytochrome P450 Activities and Drug Interactions In Vivo. Chem. Biol. 14, 1043–1051 (2007).

29. Wright, A. T., Song, J. D. & Cravatt, B. F. A suite of activity-based probes for human cytochrome P450 enzymes. J. Am. Chem. Soc. 131, 10692–10700 (2009).

30. Jean, P., Lopez-Garcia, P., Dansette, P., Mansuy, D. & Goldstein, J. L. Oxidation of tienilic acid by human yeast-expressed cytochromes P-450 2C8, 2C9, 2C18 and 2C19. Evidence that this drug is a mechanism-based inhibitor specific for cytochrome P-450 2C9. Eur. J. Biochem. 241, 797–804 (1996).

31. Donato, T., Jimenez, N., Castell, J. V & Gomez-Lechon, M. J. Fluorescence-based assays for screening nine cytochrome P450 (P450) activities in intact cells expressing individual human P450 enzymes. Pharmacology 32, 699–706 (2004).

32. Tai, G. et al. In-vitro and in-vivo effects of the CYP2C9*11 polymorphism on warfarin metabolism and dose. Pharmacogenet. Genomics 15, 475–481 (2005).

33. Matreyek, K. A., Stephany, J. J., Chiasson, M. A., Hasle, N. & Fowler, D. M. An improved platform for functional assessment of large protein libraries in mammalian cells. Nucleic Acids Res. 48, (2020).

34. P.B. Danielson, B. S. P. The Cytochrome P450 Superfamily: Biochemistry, Evolution and Drug Metabolism in Humans. Curr. Drug Metab. 3, 561–597 (2002).

35. Dickmann, L. J., Locuson, C. W., Jones, J. P. & Rettie, A. E. Differential Roles of Arg97, Asp293, and Arg108 in Enzyme Stability and Substrate Specificity of CYP2C9. Mol. Pharmacol. 65, 842–850 (2004).

36. Reynald, R. L., Sansen, S., Stout, C. D. & Johnson, E. F. Structural characterization of human cytochrome P450 2C19: Active site differences between P450s 2C8, 2C9, and 2C19. J. Biol. Chem. 287, 44581–44591 (2012).

37. Nair, P. C., McKinnon, R. A. & Miners, J. O. Cytochrome P450 structure–function: insights from molecular dynamics simulations. Drug Metabolism Reviews 48, 434–452 (2016).

38. Roberts, A. G. et al. Intramolecular heme ligation of the cytochrome P450 2C9 R108H mutant demonstrates pronounced conformational flexibility of the B-C loop region: Implications for substrate binding. Biochemistry 49, 8700–8708 (2010).

39. Johnson, E. F. & Stout, C. D. Structural diversity of eukaryotic membrane cytochrome P450s. Journal of Biological Chemistry 288, 17082–17092 (2013).

40. Gay, S. C., Roberts, A. G. & Halpert, J. R. Structural features of cytochromes P450 and ligands that affect drug metabolism as revealed by X-ray crystallography and NMR. Future Medicinal Chemistry 2, 1451–1468 (2010).

41. Williams, P. A. et al. Crystal structure of human cytochrome P450 2C9 with bound warfarin. Nature 424, 464–468 (2003).

42. Wester, M. R. et al. The structure of human cytochrome P450 2C9 complexed with flurbiprofen at 2.0-Å resolution. J. Biol. Chem. 279, 35630–35637 (2004).

43. Maekawa, K. et al. Structural Basis of Single-Nucleotide Polymorphisms in Cytochrome P450 2C9. Biochemistry 56, 5476–5480 (2017).

44. Hasemann, C. A., Kurumbail, R. G., Boddupalli, S. S., Peterson, J. A. & Deisenhofer, J. Structure and function of cytochromes P450:a comparative analysis of three crystal structures. Structure 3, 41–62 (1995).

45. Sirim, D., Widmann, M., Wagner, F. & Pleiss, J. Prediction and analysis of the modular structure of cytochrome P450 monooxygenases. BMC Struct. Biol. 10, 34 (2010).

46. Kemper, B. Structural basis for the role in protein folding of conserved proline-rich regions in cytochromes P450. Toxicol. Appl. Pharmacol. 199, 305–315 (2004).

47. Chen C. Di, Doray, B. & Kemper, B. A conserved proline-rich sequence between the N-terminal signal-anchor and catalytic domains is required for assembly of functional cytochrome P450 2C2. Arch. Biochem. Biophys. 350, 233–238 (1998).

48. Mustafa, G., Yu, X. & Wade, R.C. Structure and Dynamics of Human Drug-Metabolizing Cytochrome P450 Enzymes. in Drug Metabolism Prediction 75–102 (Wiley Blackwell, 2014). doi:10.1002/9783527673261.ch04

49. Arendse, L. B. & Blackburn, J. M. Effects of polymorphic variation on the thermostability of heterogenous populations of CYP3A4 and CYP2C9 enzymes in solution. Sci. Rep. 8, (2018).

50. Cojocaru, V., J. Winn, P. & C. Wade, R. Multiple, Ligand-dependent Routes from the Active Site of Cytochrome P450 2C9. Curr. Drug Metab. 13, 143–154 (2012).

51. Szczesna-Skorupa, E., Chen, C. I. D. I., Rogers, S. & Kemper, B. Mobility of cytochrome P450 in the endoplasmic reticulum membrane. Proc. Natl. Acad. Sci. U. S. A. 95, 14793–14798 (1998).

52. Niinuma, Y. et al. Functional characterization of 32 CYP2C9 allelic variants. Pharmacogenomics J. 14, 107–114 (2014).

53. Wang, Y. H. et al. Effect of 36 CYP2C9 variants found in the Chinese population on losartan metabolism in vitro. Xenobiotica 44, 270–275 (2014).

54. Dai, D. P. et al. In vitro functional characterization of 37 CYP2C9 allelic isoforms found in Chinese Han population. Acta Pharmacol. Sin. 34, 1449–1456 (2013).

55. Zhang, L. et al. CYP2C9 and CYP2C19: Deep Mutational Scanning and Functional Characterization of Genomic Missense Variants. Clin. Transl. Sci. 13, 727–742 (2020).

56. Correia, M. A., Sinclair, P. R. & De Matteis, F. Cytochrome P450 regulation: The interplay between its heme *and apoprotein moieties in synthesis, assembly,repair, and disposal. Drug Metab. Rev. 43, 1–26 (2011).

57. Lee, C. R., Goldstein, J. A. & Pieper, J. A. Cytochrome P450 2C9 polymorphisms: A comprehensive review of the in-vitro and human data. Pharmacogenetics 12, 251–263 (2002).

58. Rademacher, P. M., Woods, C. M., Huang, Q., Szklarz, G. D. & Nelson, S. D. Differential oxidation of two thiophene-containing regioisomers to reactive metabolites by cytochrome P450 2C9. Chem. Res. Toxicol. 25, 895–903 (2012).

59. Gietz, R. D. & Schiestl, R. H. High-efficiency yeast transformation using the LiAc/SS carrier DNA/PEG method. Nat. Protoc. 2, 31–34 (2007).

60. McDonald, M. G. et al. Expression and functional characterization of breast cancer-associated cytochrome P450 4Z1 in saccharomyces cerevisiae. Drug Metab. Dispos. 45, 1364–1371 (2017).

61. Dong, J. et al. A two-step integration method for seamless gene deletion in baker’s yeast. Anal. Biochem. 439, 30–36 (2013).

62. Sikorski, R. S. & Hieter, P. A system of shuttle vectors and yeast host strains designed for efficient manipulation of DNA in Saccharomyces cerevisiae. Genetics 122, 19–27 (1989).

63. Mumberg, D., Muller, R. & Funk, M. Regulatable promoters of saccharomyces cerevisiae: Comparison of transcriptional activity and their use for heterologous expression. Nucleic Acids Res. 22, 5767–5768 (1994).

64. Mumberg, D., Müller, R. & Funk, M. Yeast vectors for the controlled expression of heterologous proteins in different genetic backgrounds. Gene 156, 119–122 (1995).

65. Güldener, U., Heck, S., Fiedler, T., Beinhauer, J. & Hegemann, J. H. A new efficient gene disruption cassette for repeated use in budding yeast. Nucleic Acids Res. 24, 2519–2524 (1996).

66. Gibson, D. G. et al. Enzymatic assembly of DNA molecules up to several hundred kilobases. Nat. Methods 6, 343–345 (2009).

67. Jain, P. C. & Varadarajan, R. A rapid, efficient, and economical inverse polymerase chain reaction-based method for generating a site saturation mutant library. Anal. Biochem. 449, 90–98 (2014).

68. Zhang, J., Kobert, K., Flouri, T. & Stamatakis, A. PEAR: A fast and accurate Illumina Paired-End reAd mergeR. Bioinformatics 30, 614–620 (2014).

69. Rubin, A. F. et al. A statistical framework for analyzing deep mutational scanning data. Genome Biol. 18, (2017).

70. Chen, D. C., Yang, B. C. & Kuo, T. T. One-step transformation of yeast in stationary phase. Curr. Genet. 21, 83–84 (1992).

71. García-Nafría, J., Watson, J. F. & Greger, I. H. IVA cloning: A single-tube universal cloning system exploiting bacterial In Vivo Assembly. Sci. Rep. 6, (2016).

72. Pompon, D., Louerat, B., Bronine, A. & Urban, P. Yeast expression of animal and plant P450s in optimized redox environments. Methods Enzymol. 272, 51–64 (1996).

