## Supplemental Information for "Massively parallel characterization of CYP2C9 variant enzyme activity and abundance"

### SUPPLEMENTARY FIGURES

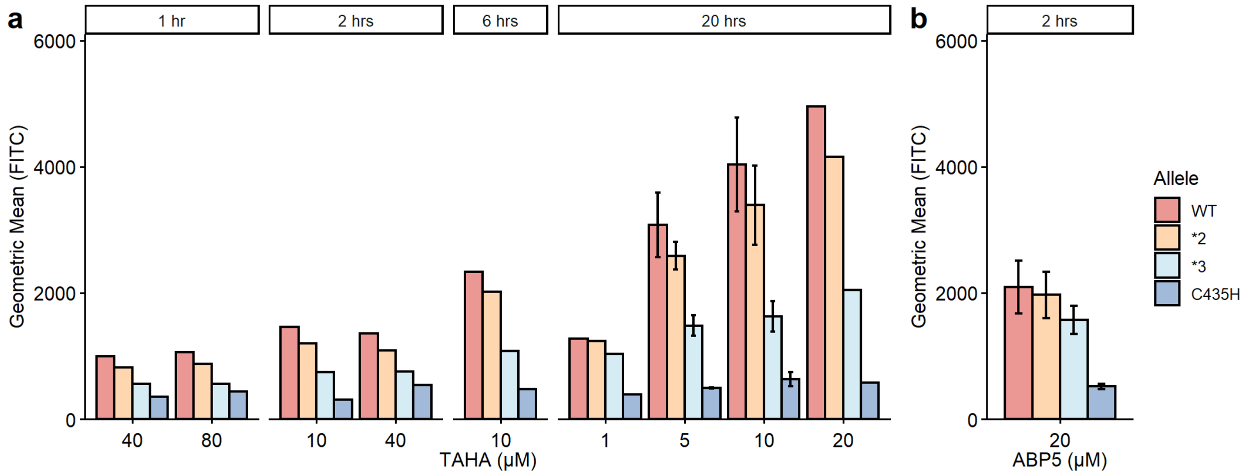


**Supplementary Figure 1. Probe labeling optimization of CYP2C9 alleles.** Barplot of flow cytometry of ABPP-labeled CYP2C9 WT (red), reduced activity alleles (*2 and *3, orange and turquoise), and null allele (C435H, blue). Cells labeled with TAHA probe (a) or ABP5 probe (b). Incubation times tested shown on top, probe concentration shown on bottom (μM). Error bars show standard deviation, each sample from ~20,000 cells.

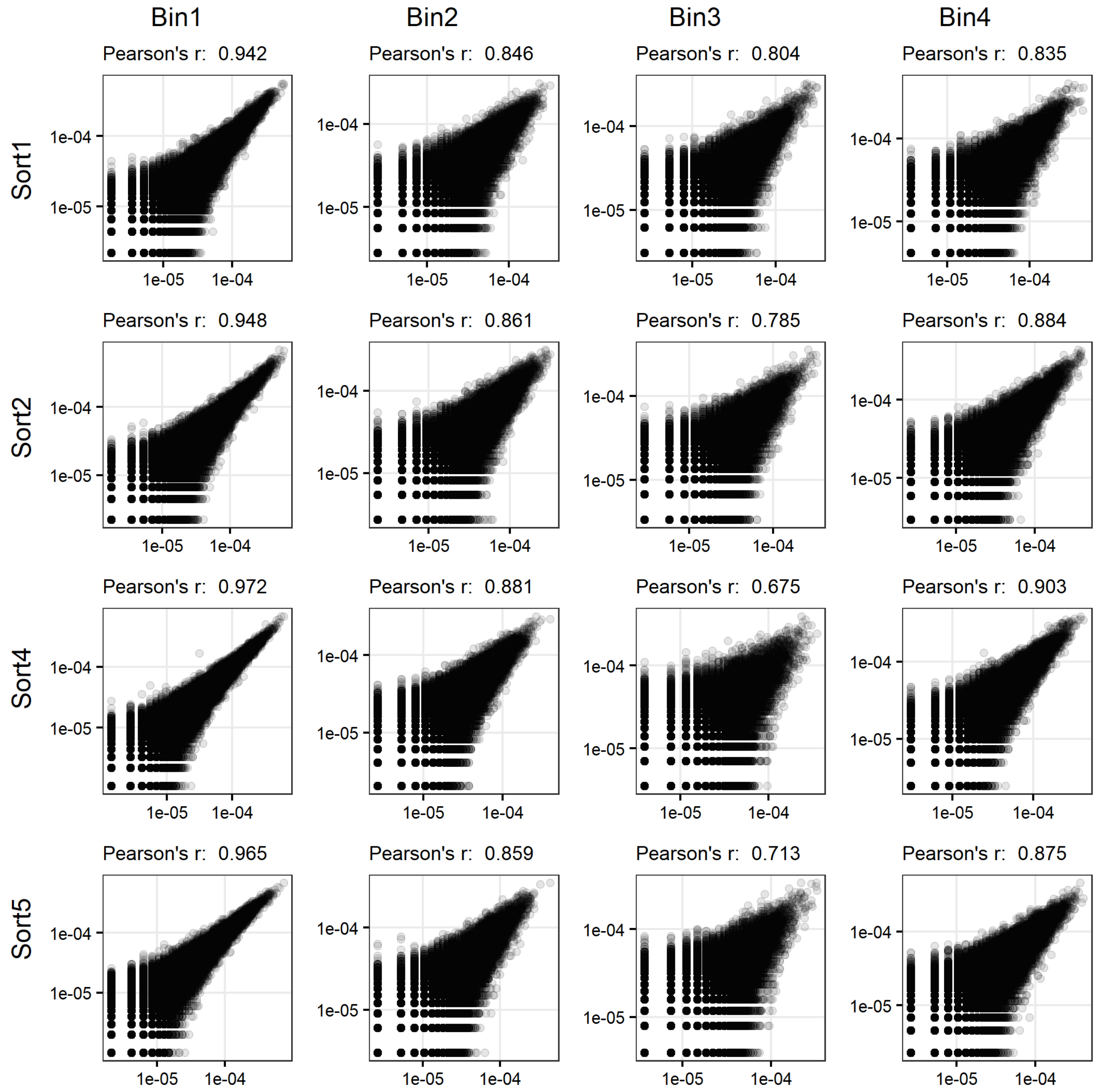


**Supplementary Figure 2. CYP2C9 activity library technical replicate correlation.** Sequencing of technical (PCR) replicates of CYP2C9 activity library: Scatterplots of barcode frequency correlation of each bin for each of the four sorts of the CYP2C9 activity library

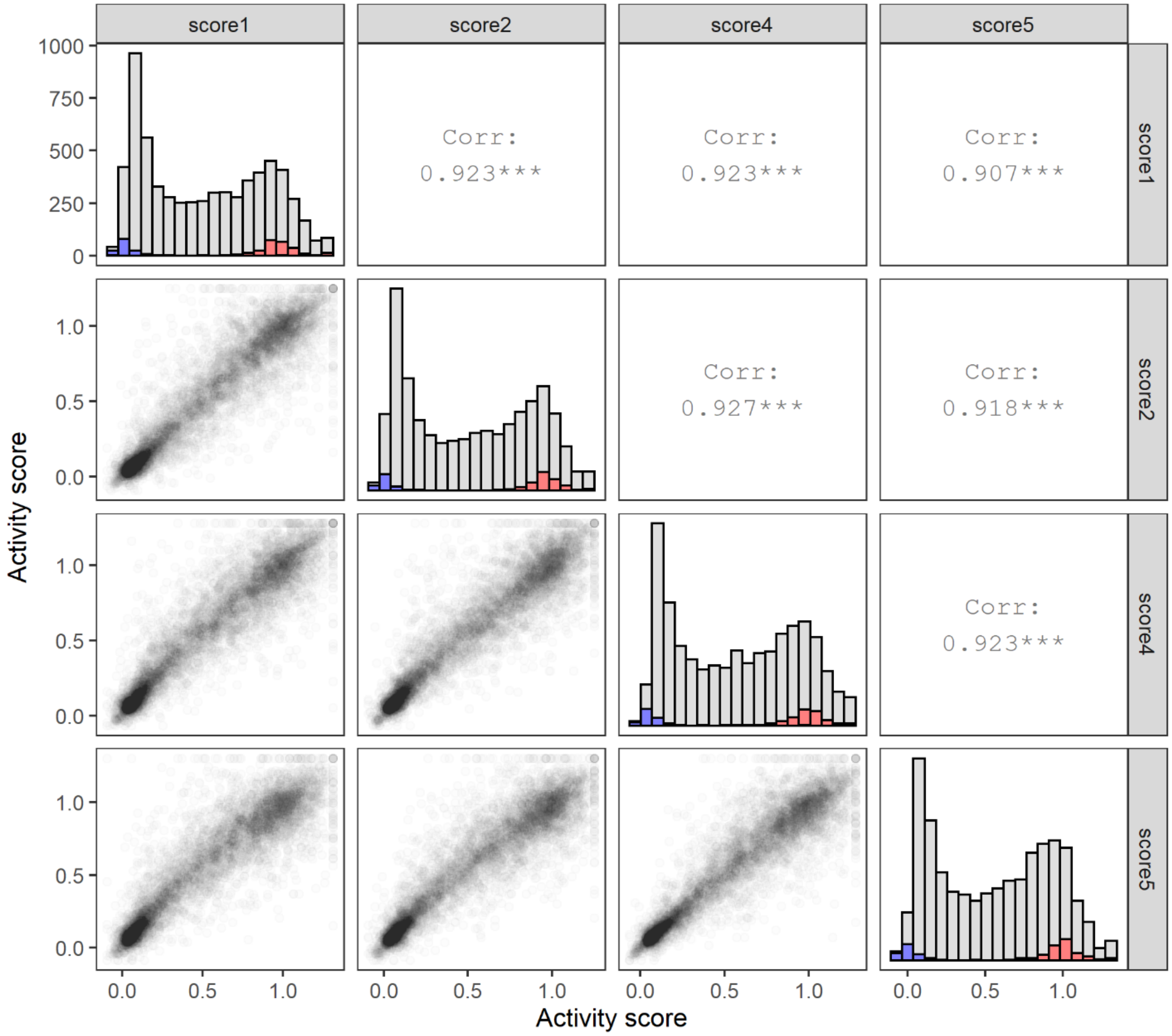

score1
score2
score4
score5
Corr:
0.923***
Corr:
0.923***
Corr:
0.907***
Corr:
0.927***
Corr:
0.918***
Corr:
0.923***
Activity score
Activity score


13

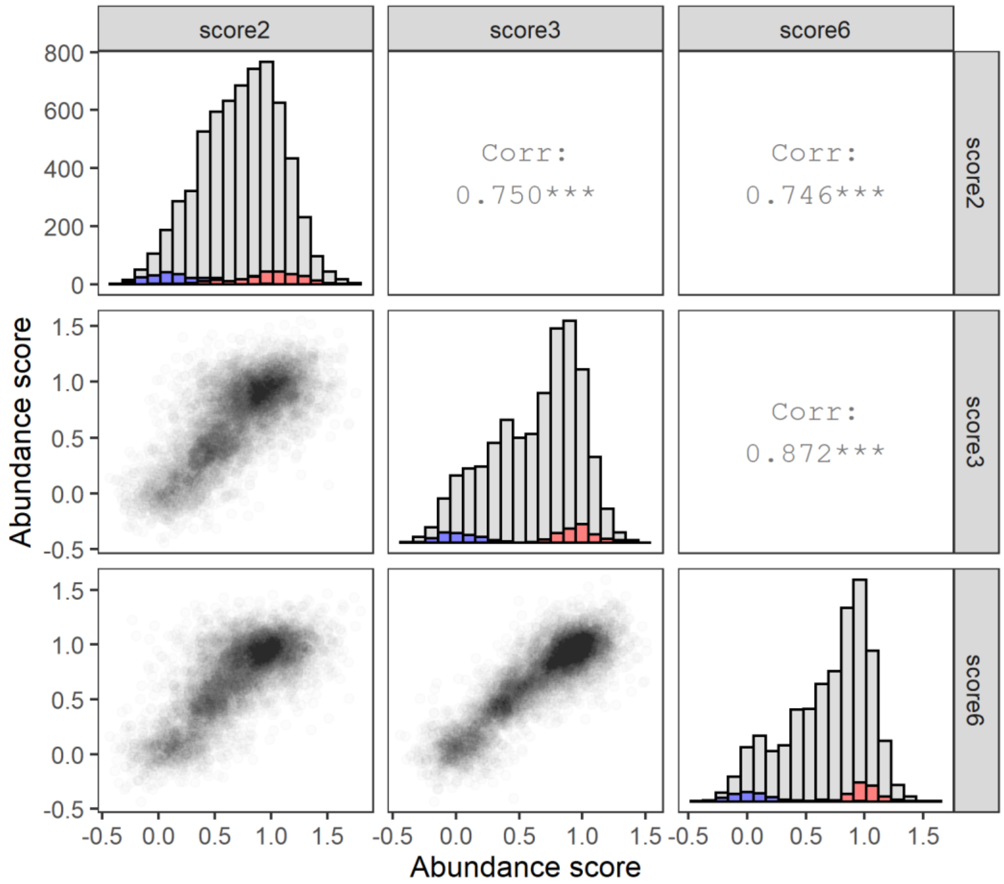


**Supplementary Figure 3. CYP2C9 score correlation matrices.** Replicate correlation of CYP2C9 activity scores for the four replicates (top), and CYP2C9 abundance scores for the three replicates (bottom). Bottom corner: pairwise scatterplot of scores, diagonal: stacked histograms of synonymous (red), missense (grey), and nonsense (blue) score distributions, top corner: Pearson's r values.

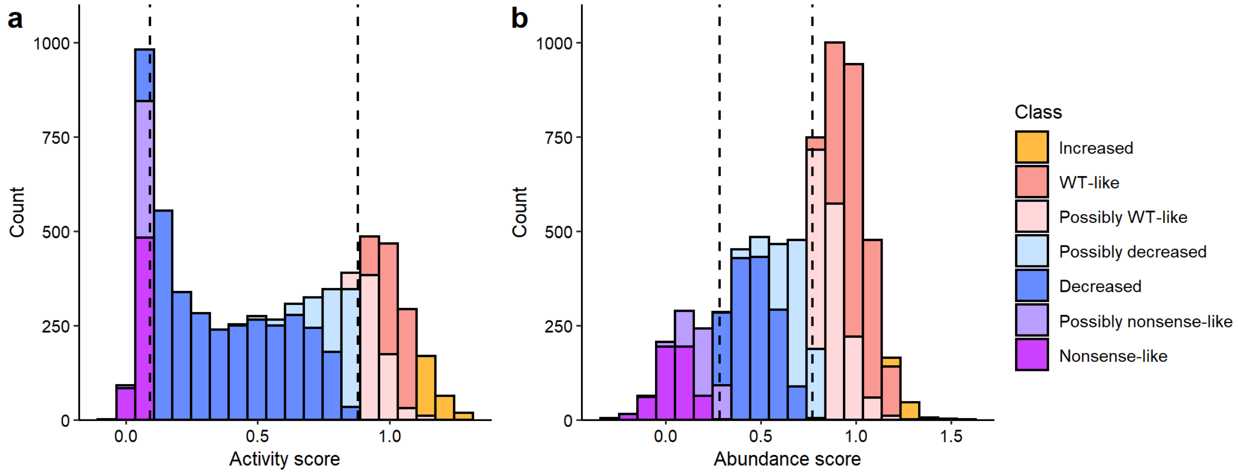


**Supplementary Figure 4. Classification of CYP2C9 scores into classes.** Stacked histograms of CYP2C9 (a) activity and (b) abundance scores categorized into classes. In dotted lines, the 95th percentile of the nonsense distribution (left), and the 5th percentile of the synonymous distribution (right), used for categorization. Variants were categorized by determining whether variant scores and confidence intervals fell within the synonymous and nonsense variant thresholds, as detailed in Methods.

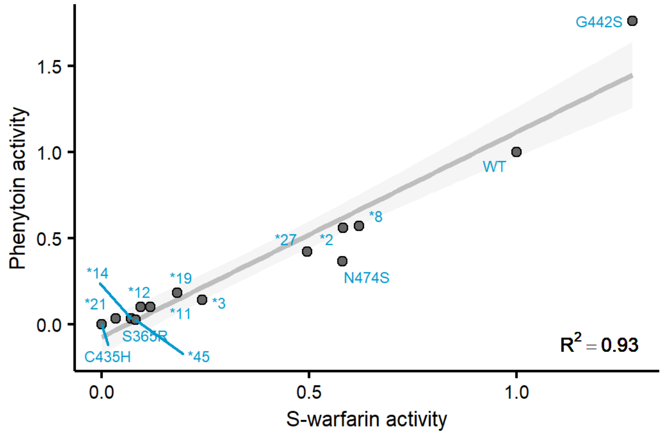


**Supplementary Figure 5. Comparison of gold-standard activity assay with different CYP2C9 substrates.** Individual CYP2C9 alleles were expressed in yeast and microsomes were extracted. The rate of S-warfarin 7-hydroxylation and phenytoin 4-hydroxylation was tested with these microsomes using LC-MS. A scatterplot comparing the CYP2C9 variant activity with these two different substrates is shown. The grey line is the regression line, and shaded area shows the 95% confidence interval. All activities are normalized to wild type rates.

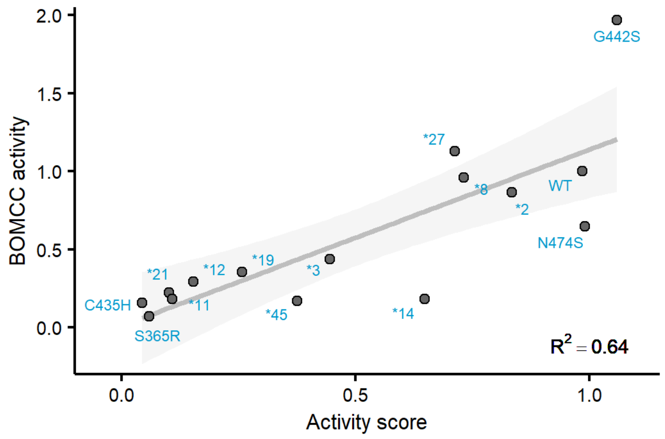


**Supplementary Figure 6. Comparison of CYP2C9 activity scores with fluorogenic activity assay in yeast microsomes.** Scatterplot of CYP2C9 Click-seq activity scores plotted against individually tested CYP2C9 alleles using a fluorogenic substrate. The conversion of BOMCC to CHC (fluorescent) by individual CYP2C9 variants was monitored using a plate reader. The grey line is the regression line, and shaded area shows the 95% confidence interval. All activities are shown normalized to wild type rates.

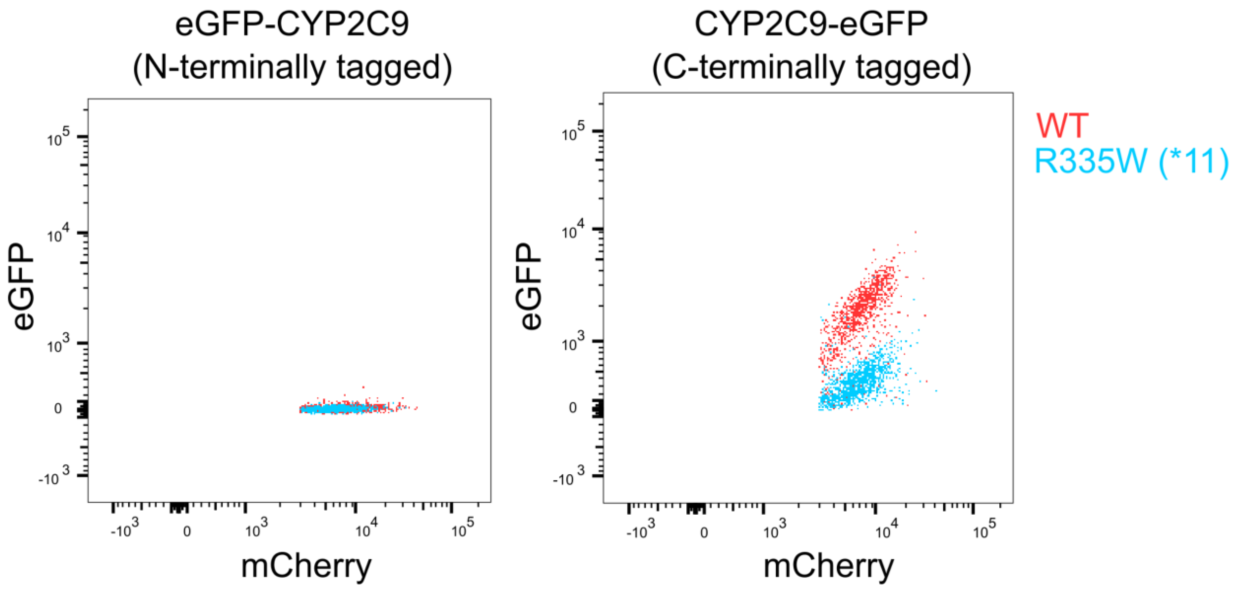


**Supplementary Figure 7. N vs. C-terminal CYP2C9 tagging.** Scatterplots of eGFP vs. mCherry fluorescence for cells expressing either N-terminally eGFP-tagged CYP2C9 (left) or C-terminally eGFP-tagged CYP2C9 (right). WT CYP2C9 shown in red, unstable R335W (*11) variant shown in blue. ~20,000 cells shown each.

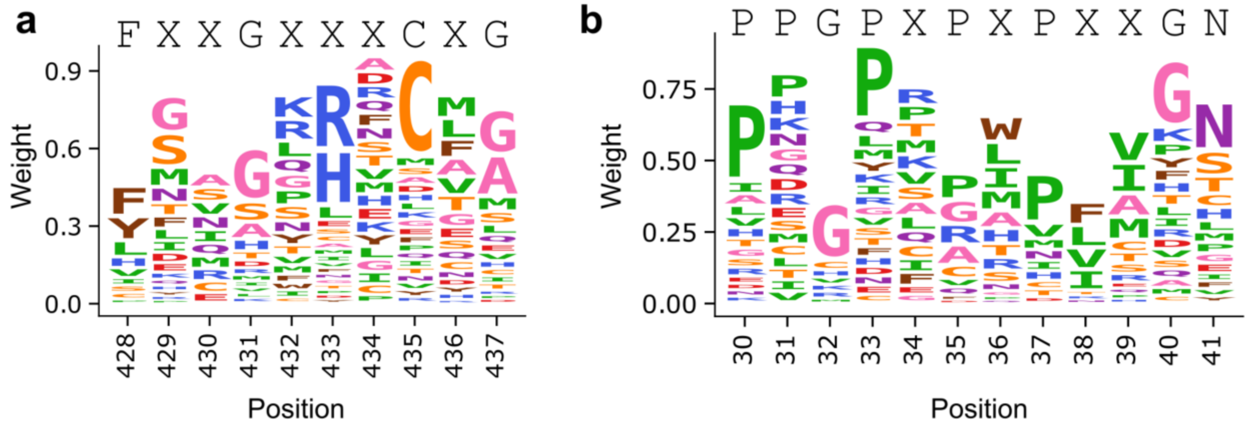


**Supplementary Figure 8. Logo plot of heme binding motif and PPGP motif from DMS data.** a) Logo plot of CYP2C9 heme binding motif using Click-seq activity scores, positions 428 to 437. Published heme binding motif[1] shown on top. b) Logo plot of CYP2C9 PPGP linker motif using Click-seq activity scores, positions 30 to 41. Published PPGP motif[2] shown on top. Variant weights calculated by rescaling activity scores from 0 to 1, calculating the frequency of each variant by position, and multiplying frequencies by the fraction of total number of variants present at each position. Variants colored by amino acid type. Figures made using dmslogo (https://github.com/jbloomlab/dmslogo).

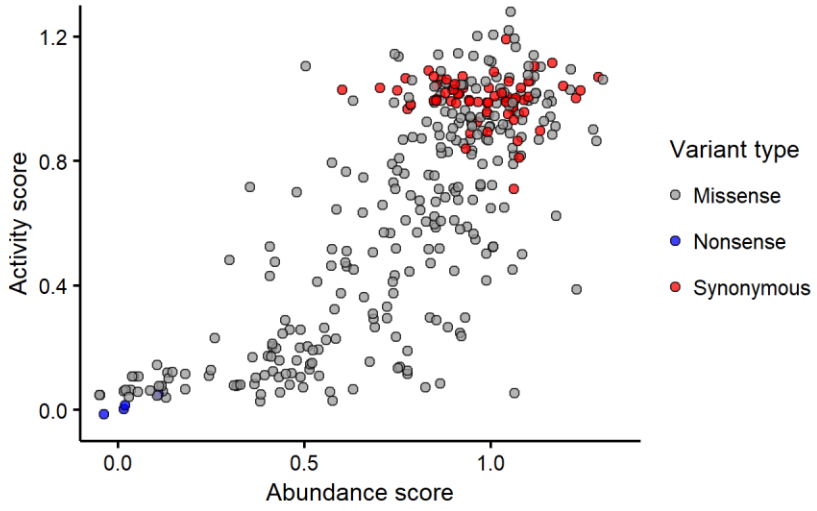


**Supplementary Figure 9. Human CYP2C9 variants with activity and abundance scores.** Scatter plot of variant activity vs. abundance score, colored by type of mutation in gnomAD. Variants combined from gnomAD v2 and v3 data, and filtered for missense, stop-gained, and missense variants. A total of 281 gnomAD variants are shown with both activity and abundance scores.

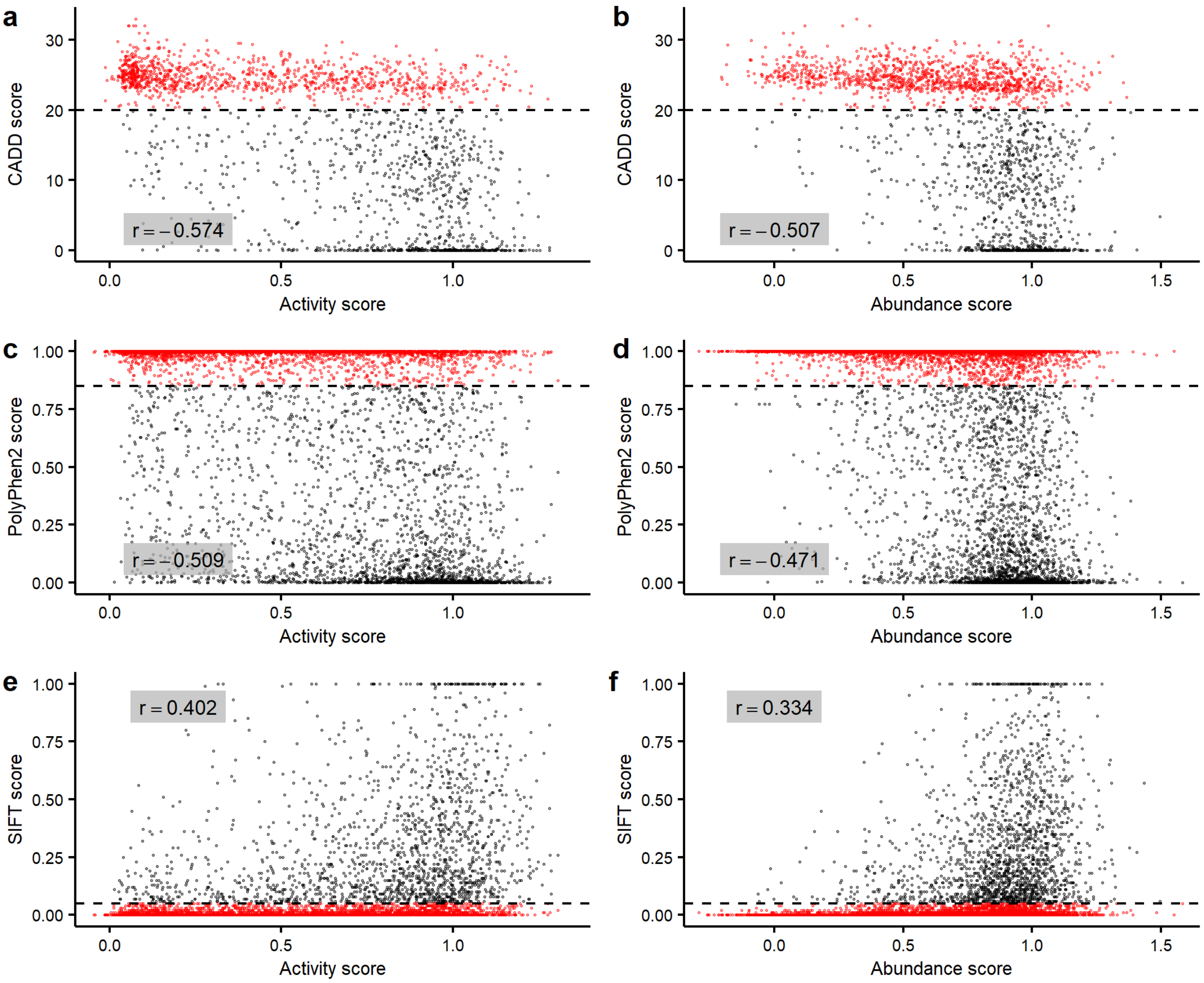


**Supplementary Figure 10. Computational prediction of CYP2C9 missense variant effect.** Scatter plots of CYP2C9 missense variant activity score (left) or abundance score (right) vs computational predictions of variant effect. For all plots, correlation (Pearson's r) shown in grey box. In a) and b), CADD score[3] vs activity or abundance score. A CADD score of >20 is considered damaging. Cutoff shown as a dotted line and points shown in red. In c) and d), PolyPhen2 score[4] vs activity or abundance score. A PolyPhen2 score of >0.85 is considered damaging. Cutoff shown as a dotted line and points shown in red. In e) and f), SIFT score[5] vs activity or abundance score. A SIFT score of <0.05 is considered damaging. Cutoff shown as a dotted line and points shown in red.

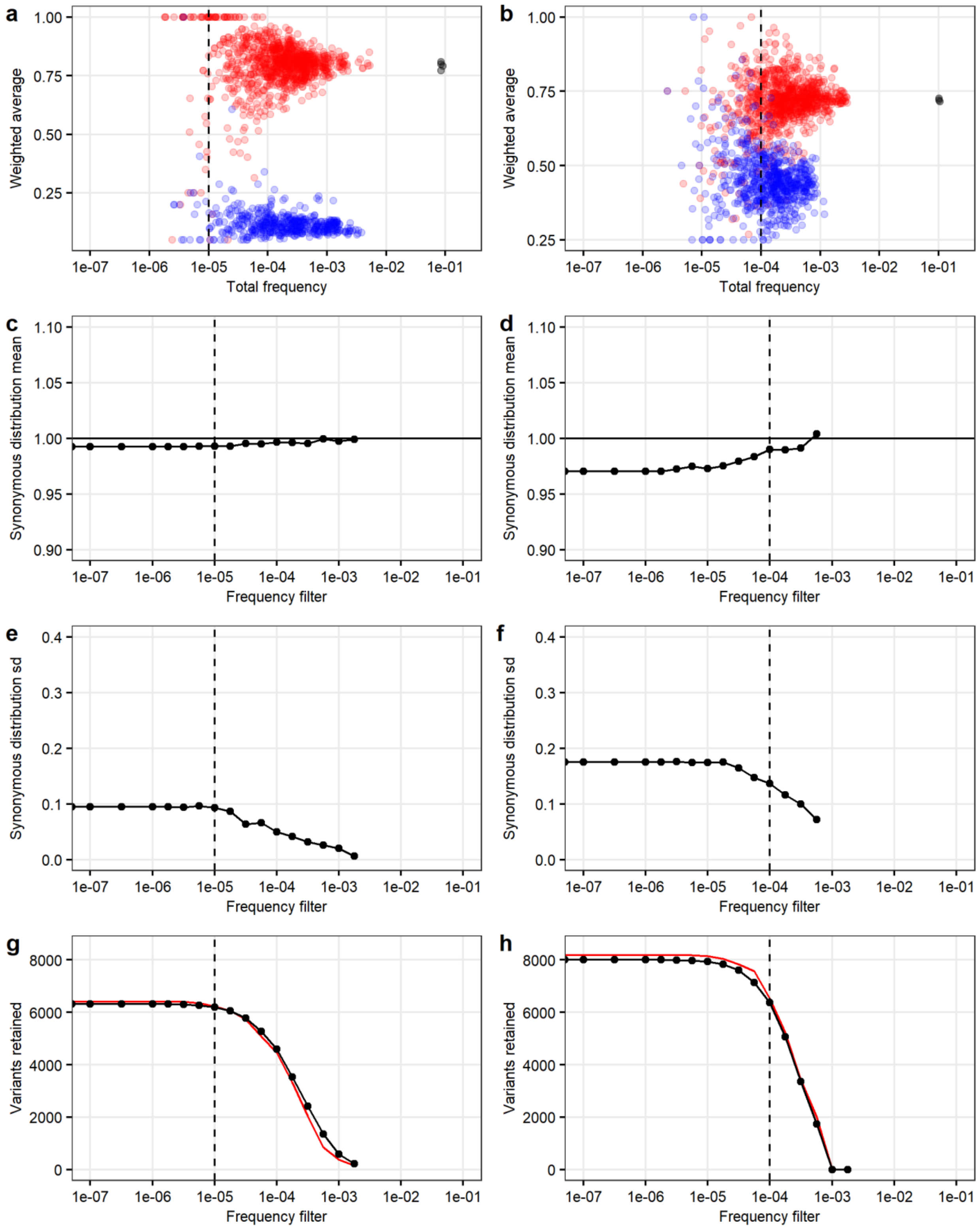


**Supplementary Figure 11. Determining variant frequency filters.** Variant frequency filter used for Click-seq in a,c,e,g) and VAMP-seq in b,d,f,h). In a and b), scatterplots of variant weighted average vs total frequency for wild type (black), synonymous (red), and nonsense (blue) variants, for each of the four or three replicates for the CYP2C9 a)

Click-seq and b) VAMP-seq libraries respectively. In c and d), the mean of the synonymous distribution at different frequency filters for the Click-seq and VAMP-seq library, respectively. In e and f), standard deviation of the synonymous distribution at varying frequency filters. In g and h), the number of missense variants retained (black) at varying frequency filters, and 25 times the number of synonymous variants retained shown in red. For all plots, the frequency filter used for library analysis is shown as a dashed line. For Click-seq this was $10^{-5}$ (left), and for VAMP-seq this was $10^{-4}$ (right).

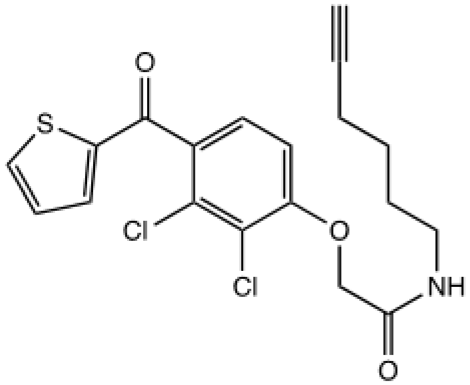


**Supplementary Figure 12. TAHA probe structure.** Chemical structure of tienilic acid hexynyl amide (TAHA).

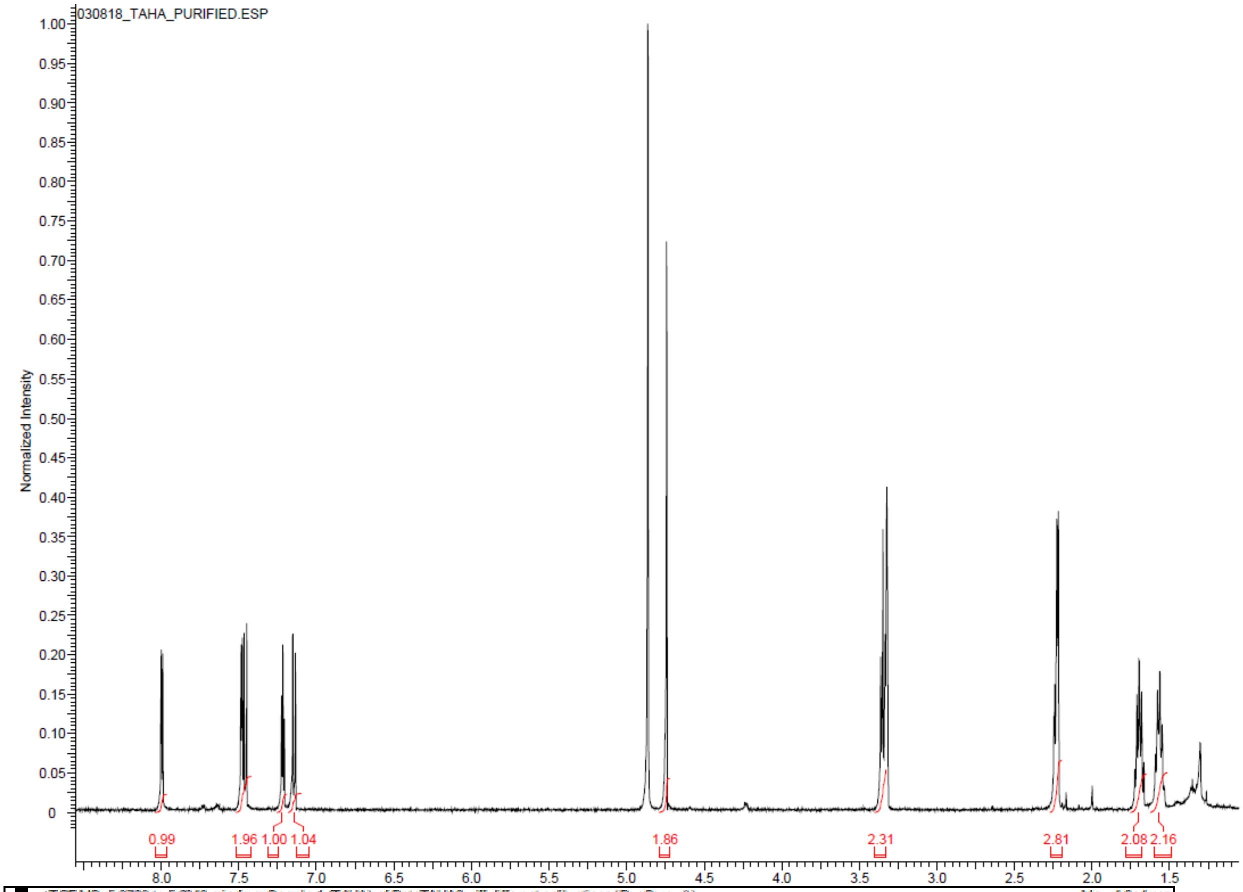
030818_TAHA_PURIFIED.ESP
Normalized Intensity
0.99
1.96
1.00
1.04
1.86
2.31
2.81
2.08
2.16

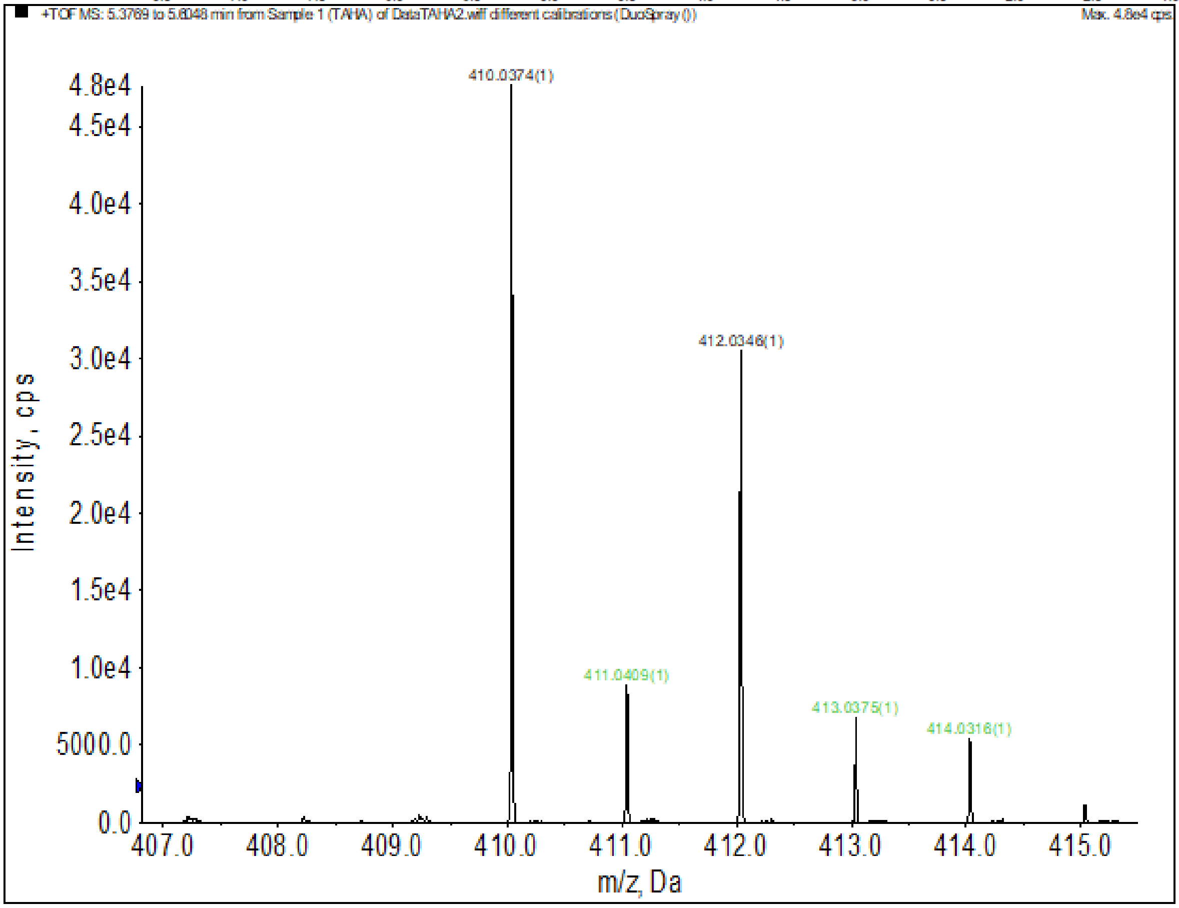
+TOF MS: 5.3769 to 5.6048 min from Sample 1 (TAHA) of DataTAHA2.wiff different calibrations (DuoSpray ())
Max. 4.8e4 cps.
410.0374(1)
412.0346(1)
411.0409(1)
413.0375(1)
414.0316(1)
Intensity, cps
m/z, Da

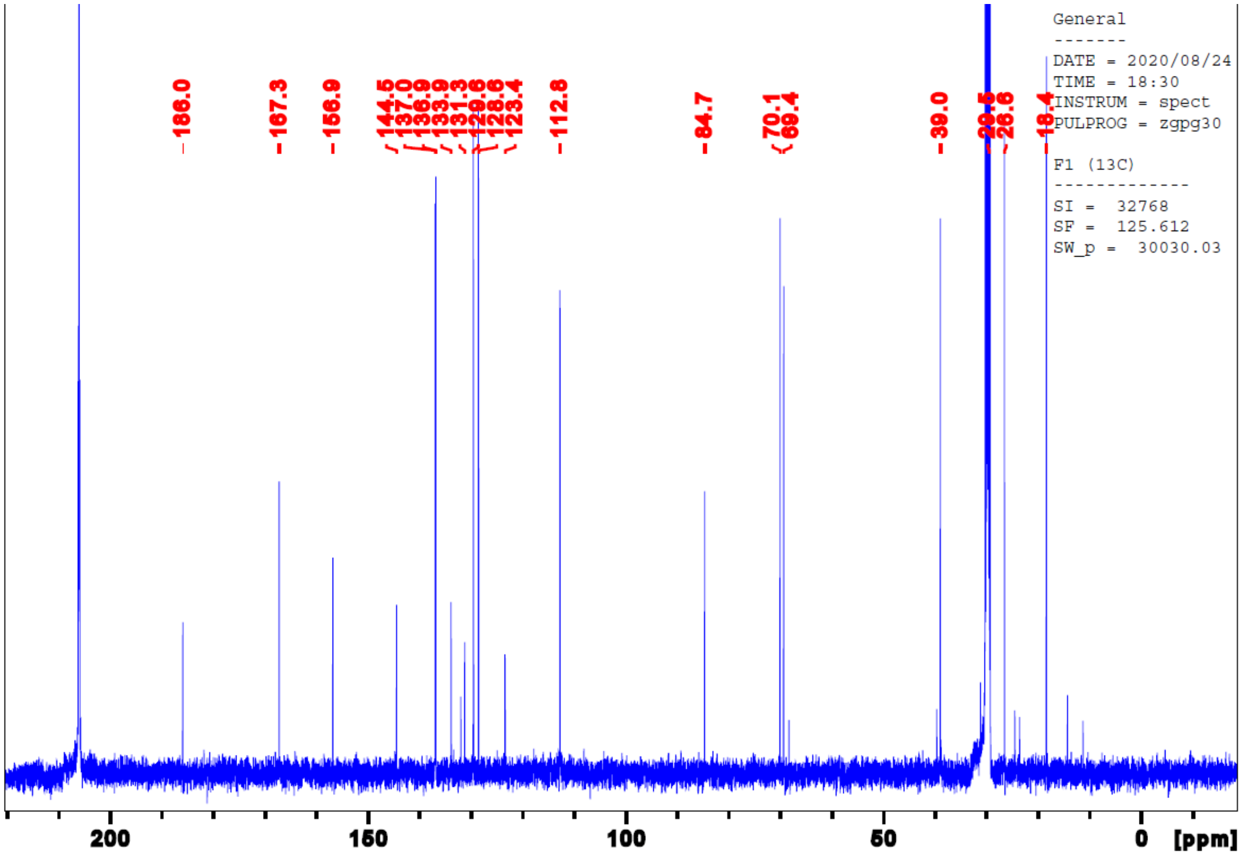


**Supplementary Figure 13. TAHA probe characterization.** (Top) $^{1}H$ spectra of TAHA , (Middle) $^{13}C$ NMR spectra of TAHA, and (Bottom) high resolution mass spectra of TAHA depicting isotopic distribution.

#### SUPPLEMENTARY TABLES

| Strain | Genotype |
|---|---|
| YMD3289 | *MATα ura3Δ0 leu2Δ1 his3Δ1 trp1Δ63 HAP1+* |
| YMD4252 | *MATa ura3Δ0 leu2- his3Δ1 trp1Δ63 HAP1+* |
| YMD4253 | *MATα ura3Δ0 leu2Δ1 his3Δ1 trp1Δ63 HAP1+ pep4Δ0 prb1Δ0* |
| YMD4254 | *MATα ura3Δ0 leu2Δ1 his3Δ1 trp1Δ63 HAP1+ pep4Δ0 prb1Δ0 HO:pGAL1-hCPR-FLAG_TRP1* |
| YMD4255 | *MATα ura3Δ0::pGPD-MYC-hb5_URA3 leu2Δ1 his3Δ1 trp1Δ63 HAP1+ pep4Δ0 prb1Δ0 HO::pGAL1-hCPR-FLAG_TRP1* |
| YMD4256 | *MATa ura3Δ0::pGPD-MYC-hb5_URA3 leu2-1 his3Δ1 trp1Δ63 HAP1+ pep4Δ0 prb1Δ0 HO::pGAL1-hCPR-FLAG_TRP1* |

**Supplementary Table 1. List of yeast strains generated in this study.**

All strains are in a *S. cerevisiae* S288C derivative background.

| Library | Experiment number | Cells sorted in Bin1 | Cells sorted in Bin2 | Cells sorted in Bin3 | Cells sorted in Bin4 |
|---|---|---|---|---|---|
| Yeast activity | 1 | 16,519,559 | 8,327,656 | 4,167,071 | 3,868,841 |
| Yeast activity | 2 | 13,747,214 | 5,500,208 | 3,056,205 | 3,165,603 |
| Yeast activity | 4 | 18,155,749 | 5,665,686 | 2,901,456 | 3,279,558 |
| Yeast activity | 5 | 15,405,689 | 5,377,714 | 3,040,153 | 3,005,108 |
| Human abundance | 2 | 625,888 | 573,840 | 624,090 | 542,210 |
| Human abundance | 3 (growout) | 1,000,000* | 1,000,000* | 1,000,000* | 1,000,000* |
| Human abundance | 6 (growout) | 1,000,000* | 1,000,000* | 1,000,000* | 1,000,000* |

**Supplementary Table 2. CYP2C9 library fluorescence activated cell sorts.** Four-way sorts of the yeast activity CYP2C9 library and the HEK 293T human abundance CYP2C9 library. For the yeast activity library, the approximate binning target percentages were: Bin1: 60%, Bin2: 20%, Bin3: 10%, Bin4: 10%. The human abundance library was binned into equal 25% bins. Unless otherwise noted in the experiment number column, DNA was amplified directly from sorted cells, rather than growing out culture and then amplifying. Asterisk indicates approximate cell numbers.

| Library | Yeast activity | Human abundance |
|---|---|---|
| SMRT cells | 2 | 2 |
| CCS reads with 10 or more passes | 309,948 | 545,277 |
| CCS reads passing filters (mapping, soft clipping, correct length barcode) | 283,195 | 515,420 |
| Unique barcodes (coverage) | 105,372 (2.9x) | 78,740 (6.9x) |
| Barcodes with one consensus read | 41,218 | 7,376 |
| Barcodes with two consensus reads | 23,806 | 8,252 |
| Barcodes with three or more consensus reads | 40,348 | 63,112 |
| Barcodes with identical consensus reads | 51,348 | 28,578 |
| Barcodes assigned with majority allele or highest quality read | Major allele: 16,906<br>Quality: 37,118 | Major allele: 42,805<br>Quality: 7,357 |
| Barcodes associated with WT CYP2C9 sequence or synonymous mutation | 2,974 | 3,697 |
| Barcodes associated with single amino acid mutation (mean, median barcodes per single amino acid mutation) | 38,127 (5.82, 3) | 49,015 (5.89, 4) |
| # single amino acid mutations (percent possible) | **6,542 (66.8%)** | **8,310 (84.8%)** |
| Barcodes associated with indel | 54,385 | 18,436 |
| Barcodes associated with two or more amino acid mutation | 9,886 | 7,592 |
| # unique nucleotide sequences | 66,958 | 37,758 |
| # unique full length nucleotide sequences | 22,421 | 22,669 |

**Supplementary Table 3. Library statistics from barcode-variant mapping.** Barcoded *CYP2C9* libraries sequenced on a Sequel II (Pacific Biosciences). CCS reads (circular consensus reads) generated with ccs2 (Pacific Biosciences).

**Additional supplementary files**

**Supplementary Table 4. Plasmids and oligos used in this study.**

**Supplementary Table 5. CYP2C9 variant activity and abundance scores.**

**Supplementary Table 6. CYP2C9 activity and abundance scores by position.**

**Supplementary Table 7. CYP2C9 Star Allele CPIC functional annotations.**

CYP2C9 star alleles and associate CPIC functional status recommendations. Star allele and associated protein variants taken from pharmvar.org. CPIC functional status and allele evidence level from[6]. CPIC functional status is biochemical functional status (normal function, decreased function, no function, uncertain function, or unknown function), and evidence level ranges from definitive (strongest), strong, moderate, limited, to inadequate evidence (weakest). Click-seq activity score, activity sd (standard deviation), and activity class (see Methods) shown.

**Supplementary Table 8. CYP2C9 individual variant validation of activity and abundance scores.**

**Supplementary Table 9. CYP2C9 optimization data with Click-seq activity-based probes.**

#### SUPPLEMENTARY REFERENCES

1. P.B. Danielson, B. S. P. The Cytochrome P450 Superfamily: Biochemistry, Evolution and Drug Metabolism in Humans. *Curr. Drug Metab.* **3**, 561–597 (2002).
2. Kemper, B. Structural basis for the role in protein folding of conserved proline-rich regions in cytochromes P450. *Toxicol. Appl. Pharmacol.* **199**, 305–315 (2004).
3. Kircher, M. *et al.* A general framework for estimating the relative pathogenicity of human genetic variants. *Nat. Genet.* **46**, 310–315 (2014).
4. Adzhubei, I. A. *et al.* A method and server for predicting damaging missense mutations. *Nat. Methods* **7**, 248–249 (2010).
5. Sim, N. L. *et al.* SIFT web server: Predicting effects of amino acid substitutions on proteins. *Nucleic Acids Res.* **40**, W452–W457 (2012).
6. Theken, K. N. *et al.* Clinical Pharmacogenetics Implementation Consortium Guideline (CPIC) for CYP2C9 and Nonsteroidal Anti-Inflammatory Drugs. *Clin. Pharmacol. Ther.* **108**, 191–200 (2020).
